# Comparative analysis of eccDNA and circRNA tools shows increased accuracy of tool combination

**DOI:** 10.1101/2025.07.14.664708

**Authors:** Aitor Zabala, Alex M. Ascensión, Iñigo Prada-Luengo, David Otaegui

## Abstract

**Introduction:** Circular nucleic acids such as extrachromosomal circular DNA (eccDNA) and circular RNA (circRNA) are increasingly recognized for their biological relevance and potential as biomarkers in disease contexts. Despite their growing importance, their detection remains challenging due to tool-specific biases, limited validation frameworks, and high variability in performance across datasets.

**Methods:** We benchmarked 10 circle detection tools across diverse conditions using both simulated and biological datasets. Our evaluation included classical performance metrics and a novel internal measure of read distribution symmetry (ΔCJ) to assess circle prediction confidence. We explored the impact of sequencing protocols, filtering strategies, and combined tool consensus.

**Results:** We found that detection accuracy was highly influenced by sequencing depth, alignment algorithm, and experimental enrichment protocols. ΔCJ proved effective in flagging potential false positive circles, showing improved accuracy of *Intersect* (circles detected by all tools) and *Rosette* (circles detected by *≥* 2 tools) combinations.

**Discussion:** This study offers a broad evaluation of circular detection tools, suggesting that the combination of *≥*3 tools is necessary for a correct prediction. These insights will inform future experimental design and data analysis pipelines in both experimental and clinical settings.

## Introduction

Extrachromosomal circular DNA (eccDNA) and circular RNA (circRNA) are covalently closed-loop structures formed via DNA circularization events and the back-splicing process of precursor messenger RNA (pre-mRNA), respectively (Cohen and Mechali, 2001; Chen and Yang, 2015). Both types of molecules are common in eukaryotic organisms and are abundant in various types of cells and tissues (Møller et al., 2015, 2018, 2020; Cohen and Mechali, 2001; Sanger et al., 1976; Memczak et al., 2013). Due to their association with various diseases, their potential use as disease biomarkers has garnered significant interest (Turner et al., 2017; Zhang et al., 2018; Iparraguirre et al., 2017).

However, eccDNA and circRNA isolation and detection poses several challenges. eccDNA and circRNAs are typically sequenced using targeted approaches, such as Circle-Seq, which is followed by nuclease treatments to eliminate linear sequences and enrichment processes to amplify circular sequences (Jeck et al., 2013; Møller et al., 2016). However, targeted sequencing is currently under debate, and alternative non-enriched techniques, including ATAC-seq and RNA-seq, are gaining significance (Iparraguirre et al., 2019; Kumar et al., 2020).

The prediction of genomic coordinates of eccDNA and circRNA circular junctions (CJ)–DNA breakpoint for eccDNA and backsplice junctions (BSJs) for circRNA–requires specialized algorithms capable of identifying reads that map to that CJ. First, reads are aligned to the reference genome and discordant reads are extracted. The unmapped reads are then remapped in reverse orientation to identify putative CJ connections. Various filtering criteria can be applied, such as the number of reads assigned to the CJ and the splicing signal that flanks the sites in the circRNA (Digby et al., 2023; Prada-Luengo et al., 2019). For the detection of circRNA, in addition to traditional methods, a pseudoreference alignment approach can also be employed. In this approach, circular read candidates are aligned with a synthetic reference that includes circular sequences to identify and validate BSJs (Salzman et al., 2012; Zhang et al., 2020).

Numerous computational software programs have been developed and tested for detecting eccDNA and circRNA. Insights from comparisons based on eccDNA (Gao et al., 2024; Li et al., 2024) and circRNA (Zeng et al., 2017; Vromman et al., 2023; Digby et al., 2024) revealed significant differences in the detection capabilities of these software, particularly in terms of the total number of circles identified. Consequently, addressing the high rate of false positives (FP) caused by technical artifacts or transcripts derived from uncommon events, such as exon duplication or trans-splicing events, remains a critical challenge (Jeck et al., 2013; Szabo and Salzman, 2016). One way to mitigate this issue is by combining two or more prediction software tools, identifying only the circles that are shared between them (Hansen, 2018). Additionally, other software works at a lower level, merging read detection results from different tools (Gaffo et al., 2021). In circRNA studies, this approach defines the so-called *bona fide* circles, increasing confidence in their detection.

Regarding the establishment of common protocols for circular analysis, *nf-core* (Ewels et al., 2020) provides a collection of community-driven high-quality Nextflow (Di Tommaso et al., 2017) pipelines for analyzing eccDNA (Schreyer et al., 2024) and circRNA (Digby et al., 2023). These pipelines can help develop standardized protocols for detecting circular molecules. Both pipelines are compatible with targeted sequencing methods as well as whole-genome sequencing (WGS) and ATAC-seq for eccDNA and RNA-seq for circRNA.

Although these tools are widely used, many studies still lack a rigorous and standardized evaluation framework, leading to considerable variability in their results. This inconsistency is largely due to the fact that most benchmarks rely solely on *in silico* data, which often produces divergent outcomes. These discrepancies arise from: (1) sequencing artifacts, (2) differences in how circles are formed in repetitive regions—leading to higher FP rates, and (3) biological post-processing steps that cannot be fully replicated computationally (Digby et al., 2024; Zhao et al., 2022). The reliance on *in silico* data is primarily due to the absence of a reliable proxy for assessing circle detection quality. As a result, most benchmarks simply compare detection outputs across tools and conditions, which is insufficient for evaluating performance on real biological data.

In this study, we present a comprehensive evaluation of five eccDNA and five circRNA detection tools using both *in silico* datasets with an array of coverages and circle sizes, as well as biological datasets produced with different methods. We show that moderate coverages (×10–×20), combined with split-read filters, minimize FP circles. To overcome the limitations of individual detection tools, we propose the *Rosette* combination, which retains only those circles supported by at least two tools, and achieves the optimal balance between precision and recall. To evaluate the accuracy of circle detection, we introduce the ΔCJ parameter—the discrepancy in read assignment to each side of the breakpoint—as a proxy for detection quality, and validate our approach on biological datasets, thereby enhancing the reliability of circular molecule quantification.

## Results

### Study design

Given the variability in individual tool performance and the differences between eccDNA and circRNA detection methods, our study evaluated five widely used tools for eccDNA detection–Circle-Map (Prada-Luengo et al., 2019), CIRCexplorer2 (Zhang et al., 2016), Circle_finder (Kumar et al., 2017), ecc_finder-bwa, and ecc_finder-minimap2 (Zhang et al., 2021)–, as well as five tools for circRNA detection–CIRCexplorer2, circRNA_finder (Westholm et al., 2014), CIRIquant (Zhang et al., 2020), find_circ (Memczak et al., 2013), and segemehl (Hoffmann et al., 2009)–. By conducting a combined evaluation, we aimed to systematically identify shared strengths and distinct limitations across these two circular molecule detection approaches.

The study consists of two separate parts making use of *in silico* and biological data. For the *in silico* analysis, we developed *CircleSim* to generate simulated data that approaches biological distributions; thus generating two separate datasets for eccDNA and circRNA with a wide range of coverages from ×5 to ×100. Reads from both *in silico* and biological data were assessed by each tool, reported circles were then filtered based on different criteria (described in Materials and Methods) and the resulting circles were then ordered based on the combinations of tools that reported them.

For *in silico* data, we evaluated a diverse range of metrics for each individual tool including (1) precision, recall, and F-score metrics, (2) variation in the coordinates reported by the tools and (3) deviations in circle detection associated to circle length.

For biological data, we analyzed Circle-Seq and ATAC-seq datasets for eccDNA, and RNA-seq datasets for an original sample–RNase(−)–and the corresponding sample after RNase R treatment–RNase(+)– for circRNA. We evaluated the same downstream filters as in the *in silico* data; as well as the tool combinations. Aside from circle detection patterns, we developed a new metric based on the discordance of reads assigned to the circular junction split site (ΔCJ) as a proxy to evaluate the “quality” of circle detection.

### Circle detection evaluation in *in silico* data

#### False positive detection is biased high coverages

Detection accuracy for eccDNA and circRNA can be significantly affected by sequencing coverage, especially given their low abundance and the specialized methods required to identify them (Tsai et al., 2017). To investigate this relationship, we evaluated how varying sequencing coverages affect the detection accuracy of eccDNA (**Figure 1A**) and circRNA (**Figure 1B**). We standardized our evaluation by defining a threshold of 20 bp to determine when two circles should be considered identical.

**Figure 1.**
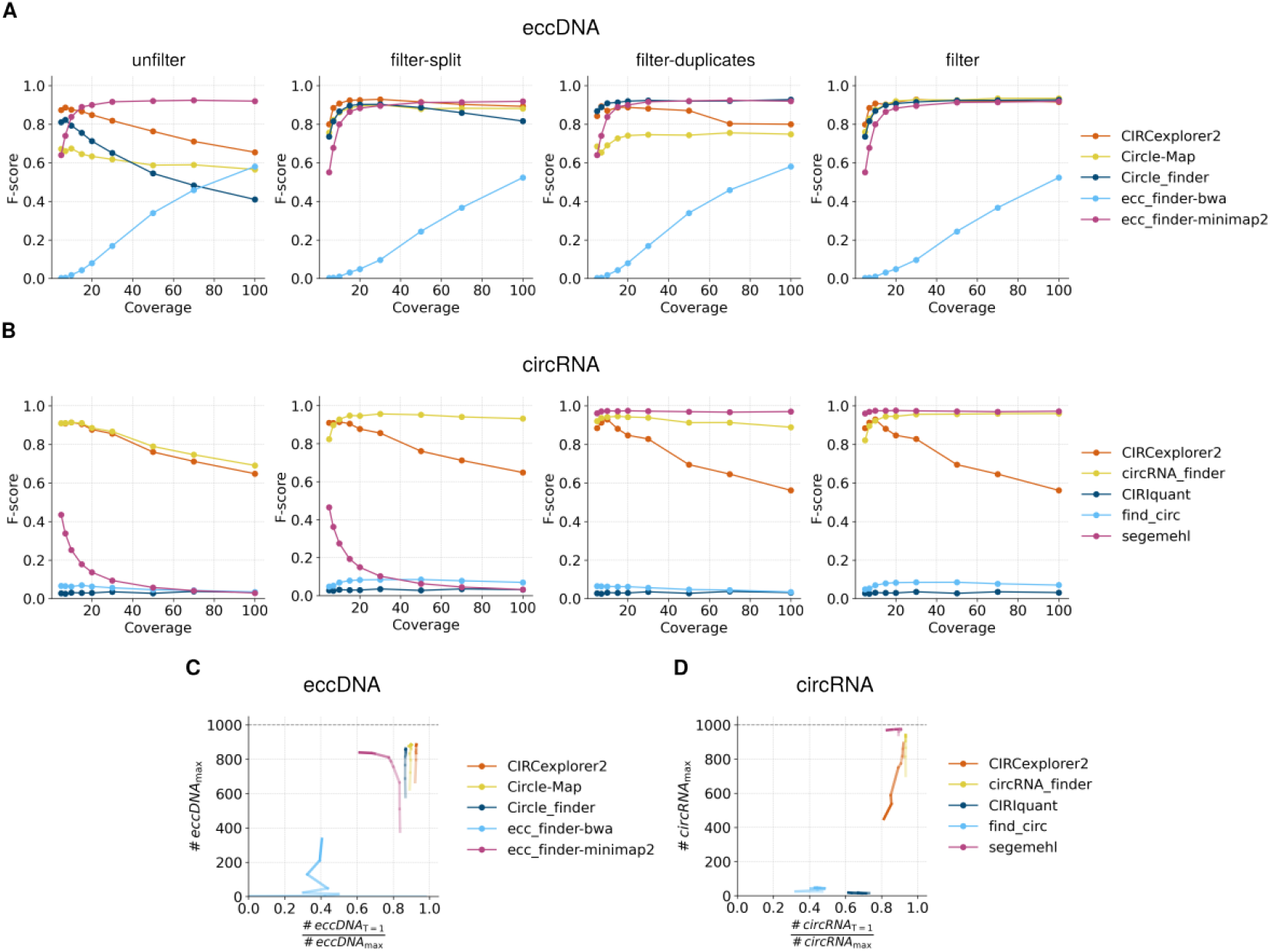
Performance analysis of detection software for eccDNA and circRNA identification in *in silico* datasets. F-score values for (**A**) eccDNA and (**B**) circRNA detection across four filtering conditions: *unfilter, filter-split, filter-duplicates*, and *filter*. Evaluation of circular predictions for (**C**) eccDNA and (**D**) circRNA based on the proportion of circles detected with an offset of 1 in the detected coordinate vs real coordinate, relative to the total number of circles detected using *filter* data. Color intensity indicates coverage level, with higher intensity corresponding to greater coverage.

Our initial results indicated that higher sequencing coverage negatively impacted overall detection performance for simulated circles, resulting in lower F-scores at higher coverage levels (**Figure 1A** and **B, *unfilter***). This decline primarily reflects a significant drop in precision, despite a modest increase in recall (**Figure S1, *unfilter*** and **Supplementary Material 1-2**). In other words, higher coverage allowed for the detection of more true positives (TP) but also substantially increased the number of false positives (FP), ultimately reducing overall accuracy.

To address this FP detection, we evaluated four filtering methods (described in Materials and Methods): (1) *unfilter*, (2) *filter-split*, (3) *filter-duplicates*, and (4) *filter*. Overall, we observed that applying any filter form increases the F-score, showing a plateau at coverages around ×10 to ×20. For good measure, the following analyses were performed using a ×30 coverage (**Table 1**).

**Table 1.** F-score values at coverage ×30 for eccDNA and circRNA detection tools under the different filtering conditions.

|  | Tool | unfilter | filter-split | filter-duplicates | filter |
| --- | --- | --- | --- | --- | --- |
| eccDNA | CIRCexplorer2 | 0.819 | 0.929 | 0.882 | 0.929 |
|  | Circle-Map | 0.618 | 0.902 | 0.746 | 0.927 |
|  | Circle_finder | 0.652 | 0.903 | 0.924 | 0.916 |
|  | ecc_finder-bwa | 0.168 | 0.096 | 0.168 | 0.096 |
|  | ecc_finder-minimap2 | 0.916 | 0.896 | 0.916 | 0.896 |
| circRNA | CIRCexplorer2 | 0.856 | 0.856 | 0.829 | 0.829 |
|  | circRNA_finder | 0.867 | 0.957 | 0.939 | 0.956 |
|  | CIRIquant | 0.035 | 0.035 | 0.035 | 0.035 |
|  | find_circ | 0.056 | 0.084 | 0.057 | 0.085 |
|  | segemehl | 0.094 | 0.103 | 0.973 | 0.974 |

Focusing on individual filter comparison, we observed that *filter-split* was generally more effective than *filter-duplicates*, although removing duplicates sometimes enhanced the effectiveness of split read-based filtering (e.g. Circle-Map in eccDNA). Among eccDNA detection tools, CIRCexplorer2 (F-score=0.929), Circle-Map (F-score=0.927), Circle_finder (F-score=0.916), and ecc_finder-minimap2 (F-score=0.896) were the most accurate (**Figure 1A, *filter***). For circRNA detection, circRNA_finder (F-score=0.956) and segemehl (F-score=0.974) were identified as the most effective software (**Figure 1B, *filter***). However, segemehl was dependent on duplicate removal (F-score=0.973) due to the high number of FPs, which were not adequately addressed by using only split read-based filtering (F-score=0.103). It is worth noting that CIRCexplorer2 output for circRNA simulated data lacked information on reads mapped to the BSJ, which diminished the effectiveness of the split read-based filtering method. Consequently, the filtering approach only managed to filter out overlapping circles and retain the longer circles. Surprisingly, find_circ (F-score=0.085) and CIRIquant (F-score=0.035) remained extremely inaccurate even after filtering. This effect is again driven by low recall values, although precision also dropped for find_circ with higher coverages; indicating that these two methods are prone to FP detection (**Figure S1** and **Supplementary Material 1-2**)

Therefore, these results show that high coverages may be unnecessary, if not detrimental for circle detection, incrementing the number of FP circles.

#### High coverage may affect the offset of circular junction detection

Accurate circular junction (CJ)–DNA breakpoint for eccDNA and BSJ for circRNA–coordinates are necessary for proper circle identification. Stemming from our hypothesis that circular coordinate accuracy may be affected by coverage, we analyzed how stable circle identification was with different coverages. To do this, we calculated the ratio of circles detected with an offset of 1 to the total number of detected circles using *filter* (**Figures 1C** and **D**). Ratios approaching 1 indicate that most circles are accurately captured with little-to-no offset, while lower ratios indicate that higher offsets are required, and therefore circle detection is less accurate.

In eccDNA we observed that for CIRCexplorer2, Circle-Map and Circle_finder the ratio was not affected by coverage, and remained at around 0.9. On the other hand, the ratio decreased with coverage for ecc_finder-minimap2, indicating that, although more circles were detected, even if those circles were “correct” based on the F-score, they were detected with a higher offset from the coordinate of the simulated circle, indicating that CJ sequence was affected by some tool-related factor. Lastly, ecc_finder-bwa showed a “stabilization” of the ratio at around 0.4 with increasing coverage, but the ratio is still low.

Regarding circRNA, two ratio trends arise. On the one hand, tools with low overall detection (find_circ and CIRIquant) show reduced ratios; whereas tools with high accuracy (segemehl and circRNA_finder) show higher ratios. Interestingly, CIRCexplorer2, which showed a decrease in circle detection accuracy with increased coverage, shows also a decreased ratio.

Therefore, it is clear for both eccDNA and circRNA that tools with low F-scores tend to show low 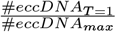 ratios showcasing that for tools with higher FP detection values, detected circles are also inaccurately detected, with higher coordinate offsets than their counterparts.

#### Specific tools showed a circle length anomaly for short circles

After observing that circle detection accuracy was variable across tools, we were interested in testing biases in the detection of circles of specific lengths. Theoretically, since *in silico* circles are generated at random positions in the genome, circle length should not pose a bias in their detection. Expectedly, tools with good circle detection accuracy showed length distributions similar to the expected distribution; whereas for tools with lower accuracies the detection is equally reduced across length (**Figure 2** and **Figure S2, left**). This effect was present both for eccDNA (ecc_finder-bwa, Kolmogorov-Smirnov test, KS=0.55 and *p*=3.97*·*10^−3^) and circRNA (CIRIquant, KS=0.75 and *p*=9.55*·*10^−6^; find_circ, KS=0.60 and *p*=1.12*·*10^−3^).

**Figure 2.**
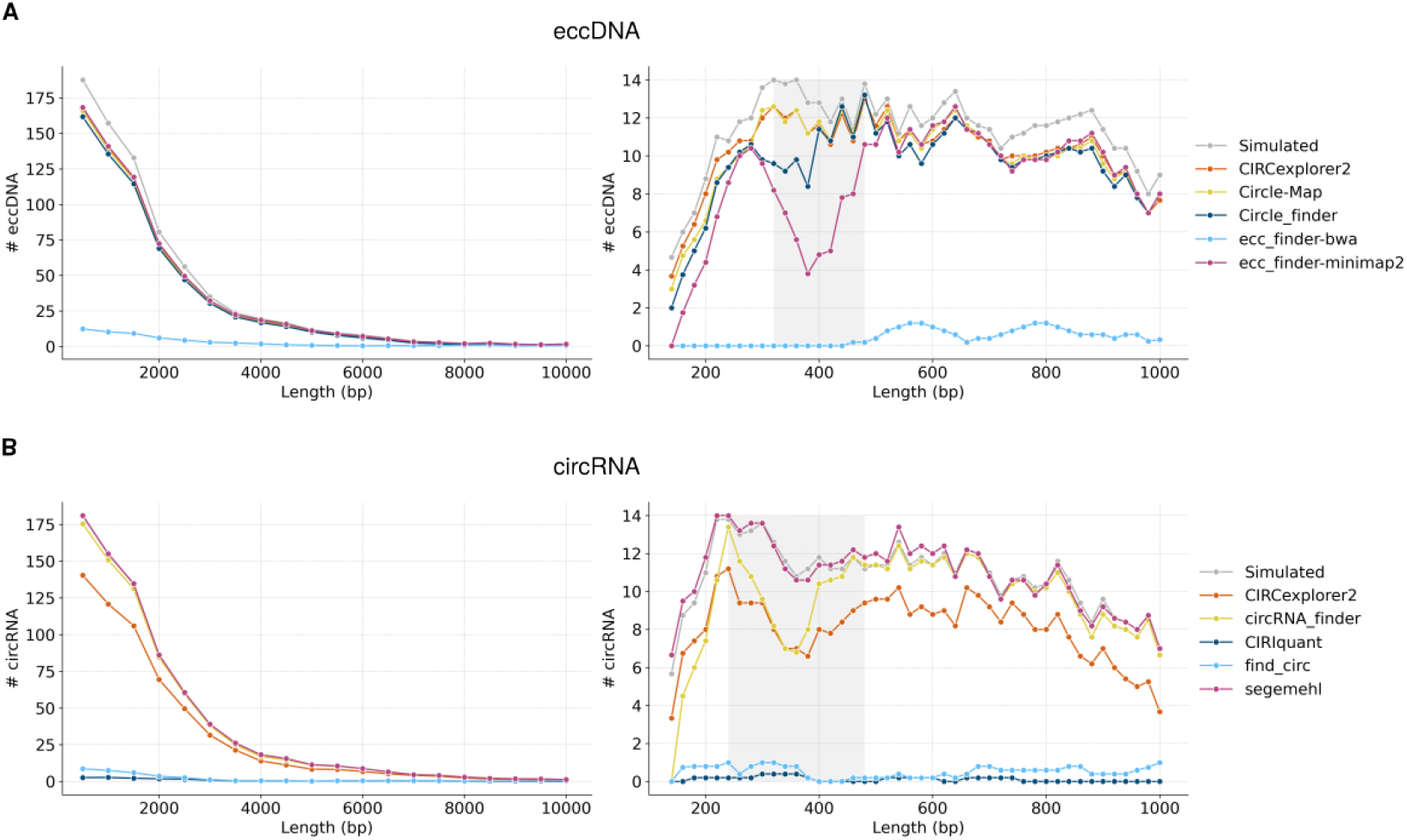
Circular length distribution in *in silico* datasets. Length distribution of detected (**A**) eccDNA and (**B**) circRNA across all size ranges (**left**) and within the short-length range (**right**) in *in silico* datasets. A grey-shaded area highlights the length interval where detection performance was lowest (eccDNA: 320-480 bp; circRNA: 240-480 bp). To enhance the clarity of the distribution plots, a sliding window of size 5 was applied to smooth the distribution curves.

Considering that the original circle distribution is condensed in shorter circles, we performed an additional analysis on short circles, ranging from 175 to 1,000 bp. Interestingly, a distinct reduction of detected circles occurred in the range of 320 to 480 bp for eccDNA (**Figure 2A** and **Figure S2A, right**) and 240 to 480 bp for circRNA (**Figure 2B** and **Figure S2B, right**). Although this reduction was more apparent for tools that already show a low circle detection, some tools with good performance showed much lower than expected counts near the 400bp mark in eccDNA Circle_finder (KS=0.38, and *p*=0.66) and ecc_finde-minimap2 (KS=0.75, and *p*=1.87*·*10^−2^) in eccDNA; and CIRCexplorer2 (KS=0.58, and *p*=3.14*·*10^−2^) and circRNA_finder (KS=0.33, and *p*=0.54) in circRNA (**Supplementary Material 3**).

#### Tools showed a biased detection of specific repeated and genomic elements

The genome is not uniform across its sequence, having areas with specific repetitive sequences that, we hypothesize, may affect the ability to detect circles. We are also interested, specially for circRNAs, if there are notable differences in the detection depending on the genomic elements they originated from.

Across the different genomic regions, the repeat element analysis showed the greatest limitation in the detection of satellite eccDNA. Among the 32 eccDNAs laying in satellite regions, all tools showed a lack of accuracy in detecting these circles (F-score ≈ 0.2, **Figure 3A, left**). These low scores were mostly driven by a lack of recall, although 3/5 tools showed reduced precision values (0.4 - 0.7) showing that FP circles were also assigned to satellite regions (**Figure S3A, left**). For the rest of elements the F-scores were generally high, with moderate decreases in regions without a demarcated repeating element (Ø).

**Figure 3.**
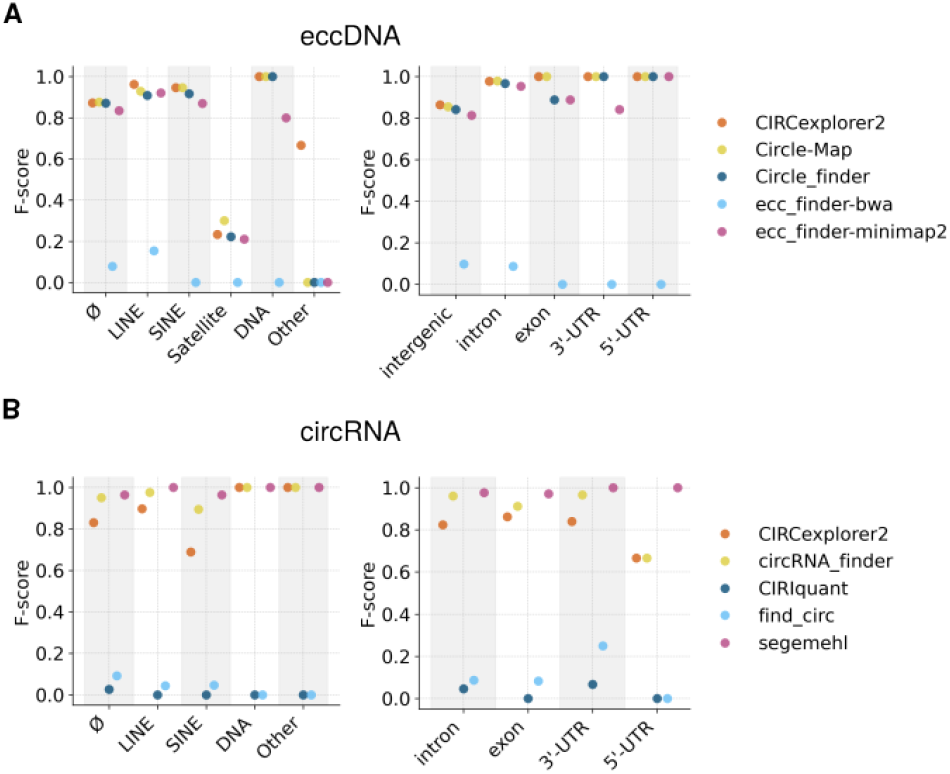
Repeat and genomic element analysis in *in silico* datasets. F-score values of repeat elements (**left**) and genomic features (**right**) associated with detected (**A**) eccDNA and (**B**) circRNA in simulated datasets. In circRNAs no intergenic circles were generated and thus the region is not included in this plot.

There was a marked decrease in accuracy for circles detected in “other–repetitive elements not marked within the SINE, LINE and similar categories– however, it is important to note that only two eccDNAs were generated in these regions, meaning that conclusions should be drawn with caution (**Supplementary Material 4**).

In the case of circRNA, no satellite regions were created since they were generated from the transcriptome. Interestingly, CIRCexplorer2 and circRNA_finder exhibited a decrease in their original F-score in SINE regions (**Figure 3B** and **Figure S3B, left**; and **Supplementary Material 5**), although this does not happen with eccDNAs.

The analysis of genomic elements showed a considerable decrease in the F-score for eccDNA in intergenic regions (F-score ≈ 0.8), due to a low recall. This suggests a significant limitation in the detection of eccDNA in these areas (**Figure 3A** and **Figure S3A, right**; and **Supplementary Material 6**). In contrast, circRNA showed a decrease in the F-score specifically in the 5’UTR regions. However, it is important to note that, in this case, only two circRNAs were generated in these regions, so the conclusions should be interpreted with caution (**Figure 3B** and **Figure S3B, right**; and **Supplementary Material 7**).

#### Combination of 3 or more tools improved circle detection in in silico data

Although individual tools may perform correctly in specific analyses, we hypothesise that the combination of several tools may improve the accuracy of circle detection, leveraging the strengths and reducing individual tool biases. This is particularly necessary for biological data, where the ground truth is unknown and circle detection may be more difficult to perform due to unknown variation in their sequences.

Although this claim is already supported in the literature, mentioned as *bona fide* circles (Hansen, 2018), these circles are usually defined by the combination of two tools. Our aim is to extend it by including three or more tools, under the assumption that the inclusion of more tools may increase the credibility of the detection of common circles. We evaluated the performance of software combinations using five different strategies (described in the Materials and Methods): *Union, Rosette, Intersect, Double*, and *Unique*. Briefly, *Union* is the set of all circles; *Instersect* is of circles detected by all tools while *Unique* refers to circles detected by one tool. *Rosette* is defined as the set of circles detected by two or more tools; whereas *Double* includes circles detected by two or more tools, excluding *Intersect* circles. *Double* circles are used as a comparison to *Rosette*, to see the effect of a more lenient detection of circles. Each strategy is evaluated with 16 tool combinations across the four filtering methods.

Two of the worst combination strategies were *Intersect* and *Unique*, both in eccDNA (**Supplementary Material 8-11**) and circRNA (**Supplementary Material 13-16**) (**Figure 4**). The low F-scores were explained by a low precision and recall in *Unique*, and by a low recall but high precision in *Intersect. Unique* circles show a low precision and recall values, indicating that simulated (true) circles are not likely to be detected by one tool (low recall) and, also, a *Unique* circle is likely not to be a true circle (low precision).

**Figure 4.**
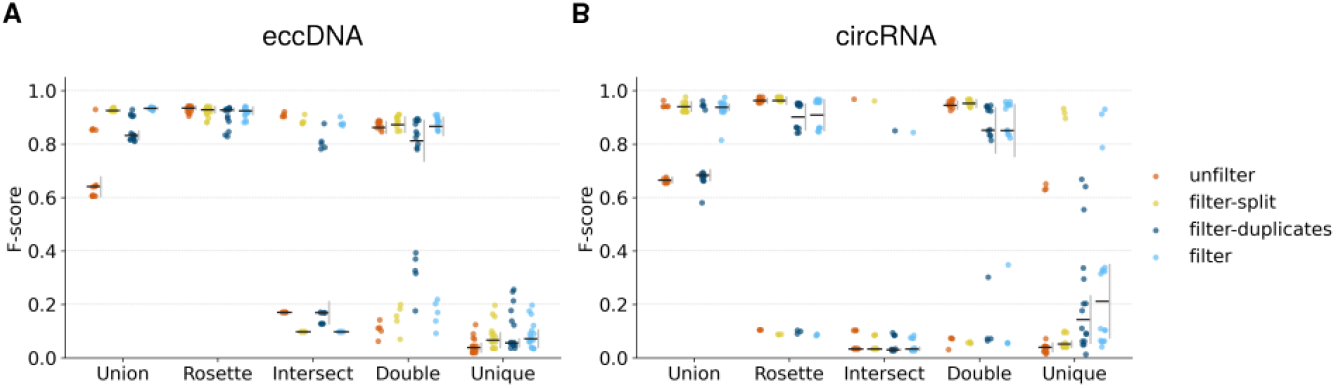
Performance analysis of software combinations for eccDNA and circRNA identification in *in insilico* datasets. Strip plot of the F-score of software combination strategies—*Union, Rosette, Intersect, Double*, and *Unique*—were evaluated under four filtering conditions: *unfilter, filter-split, filter-duplicates*, and *filter*. F-score values are shown for (**A**) eccDNA and (**B**) circRNA. For each combination-filter pair, the horizontal bar represents the mean value and the vertical gray bar represents the standard deviation.

Nonetheless, for some tool combinations–the best performing ones–*Intersect* yields high precision values. In fact, we see a bimodal distribution for recall, indicating that some tool combinations detect the simulated circles more accurately; although this detection is thus highly dependent on the combination of tools. This effect will be later discussed for biological data.

Regarding *Union* strategy, where all circles detected by all tools are included, it showed high F-scores (>0.8 for most combinations), although their precision values were low, especially for *unfilter* circles. This effect may be driven by the inclusion of *Unique* circles, which already are linked to low F-scores. However, this showcases that a proper filtering may greatly improve the detection quality.

Lastly, the two main strategies to discuss are *Rosette* and *Double*. Both strategies offer a balance between precision and recall. To perform these comparisons quantitatively, we first discuss the effect of the filtering method in the results. As expected, filtering resulted in slightly lower recall values (more false negatives), and improved precision values, specially for *Union* and *Unique* methods. However, this effect was less pronounced for *filter-duplicates* than for *filter-split*. Thus, F-scores were overall higher in *filter-split* than *filter-duplicates*; and sometimes were similar to *unfilter* or *filter* depending con the combination strategy.

Focusing on *filter-split*, for eccDNA, *Rosette* yielded higher F-scores than *Double* (two-tailed Dunn’s test, *p*=0.016) as a result of a lower, although not significant, recall (*p*=0.253). This difference was visible for the rest of filtering strategies, where *Rosette* showed a similar if not higher recall (**Supplementary Material 12**). For circRNA, *Rosette* and *Double* showed similar results. In fact, both F-scores, precision and recall values were non significant (*p*=1 for all three) (**Supplementary Material 17**).

Overall, these findings highlight that the combination of three or more tools applied by *Rosette* improves detection accuracy for both eccDNA and circRNA. The choice between methods depends on whether maximizing precision, recall, or if a balanced performance is the priority. Additionally, filter option may affect the quality of the detection, with *filter-split* showing better results than the rest of filtering options.

### Circle detection evaluation in biological data

Analysis of eccDNA and circRNA in biological data were performed with datasets with different processing techniques. For eccDNA, datasets generated using Circle-Seq and ATAC-seq were used; and for circRNA a dataset without RNAse enrichment–RNAse(−)–and with RNAse enrichment–RNAse(+)–were used. Each method has specific particularities affecting the quantity and quality of circle detection that will be further down discussed.

#### Combined tool usage and filter-split improved eccDNA and circRNA detection

To better understand the detection dynamics among circular detection tools, filtering strategies, and circle enrichment techniques, **Figures S5-S8** show, for each method, the number of circles detected by each combination of tools (**Supplementary Material 18**).

In eccDNA, we observed higher detection rates using ATAC-seq compared to Circle-Seq. For instance, for the *unfilter* case, ATAC-seq detected 67,930 circles (5528.60 circles per million of reads) whereas Circle-Seq detected 15,498 circles (1208.04 circles per million of reads). However, Circle-Seq exhibited a higher proportion of circles retained by multiple tools–e.g., 28.0% (689/1,474) retained when filtered for detection by three or more tools, compared to 8.94% (2,032/22,713) in ATAC-seq–. Although the total amount remains higher in ATAC-seq, Circle-Seq proportionally showed more inner consistence in circle detection.

Furthermore, the number of circles detected by four of five tools is extremely low in ATAC-seq (3/67,390 = 0.004% unfiltered circles), while Circle-Seq had a significantly higher ratio (642/15,489 = 4.14%) (*χ*^2^ test with Yates correction, *χ*^2^ = 2790.67, *p* < 0.0001). These observations suggest that Circle-seq provides stronger evidence for the detection of potentially higher-quality circles.

Regarding filtering strategies, both for ATAC-seq and Circle-Seq, *filter-split* retained fewer circles compared to *filter-duplicates* (36.7% vs. 74.2% for ATAC-seq and 23.02% vs. 31.6% for Circle-Seq). Nevertheless, the proportion of circles detected by three or more tools is considerably higher using *filter-split* (12.9% vs. 3.07% in ATAC-seq and 32.0% vs. 5.62% in Circle-Seq). Assuming that circles detected by multiple tools represent higher confidence detections, these results suggest that *filter-split* selects circles of higher confidence compared to *filter-duplicates*.

In terms of tool-specific performance for ATAC-seq, we identified tool groups with high joint detection (**Figure S5**): Circle_finder combined with CIRCexplorer2, Circle-Map individually, and ecc_finder-minimap2 combined with ecc_finder-bwa. These groups persist regardless of the filtering strategy, although circles detected exclusively by Circle-Map decrease when applying *filter-duplicates* and *filter*. In Circle-Seq, the groups ecc_finder-minimap2 combined with ecc_finder-bwa, Circle_finder combined with CIRCexplorer2, and CIRCexplorer2 individually appear (**Figure S6**). However, only ecc_finder-minimap2 combined with ecc_finder-bwa remains robust across different filters. This indicates clear similarities in circle detection among specific tool groups.

Similar dynamics were observed in circRNA detection. Initially, RNase(+) reported a higher number of detected circles compared to RNase(−). For the *unfilter* case, in RNAse(+) 51,909 circles were detected (2208.42 circles per million of reads) whereas in RNase(−) 15,653 circles were detected (438.64 circles per million of reads). However, a lower proportion of circles subsequently passed the filters (e.g. *filter-split* : 11,139 (21.4%) for RNase(+) vs. 6,901 (44.1%) for RNase(−)). This suggests that RNAse enrichment might increase sensitivity, requiring a more stringent filtering.

Furthermore, the filtering effect observed for circRNA mirrors that observed for eccDNA, with *filter-split* yielding a slightly higher percentage of consensus circles compared to *filter-duplicates* (RNase(+): 35.6% (3,969/11,139) vs. 22.8% (7,327/32,110); RNase(−): 30.5% (2,106/6,901) vs. 25.3% (2,392/9,469)). This reaffirms that, when choosing one strategy, *filter-split* is the superior filtering strategy. However, applying both filtering strategies yields a similar net number of circles (RNase(+) 5,538/51,909 (8.1%) vs. RNase(−): 4,620/15,653 (25.5%)), indicating that a two-filtering system, if possible, may stabilise the number of circles.

#### ΔCJ is a proxy measure of circle detection quality

One of the limitations of biological data in this context is that no ground truth is available, and thus alternative metrics must be employed to assess detection performance. In this study, we propose the difference in read coverage across each nucleotide of the CJ as a proxy for precision. The rationale behind this metric is that, assuming that reads mapped to the CJ are detected on the left side of the CJ at the same rate as on the right side of the CJ, any imbalance in the detection (e.g. a circle with 2 reads assigned on the left side and 28 on the right side) is likely to be a result of an incorrect circle detection; for which a better suited circle may or may not be available.

To show examples of this rationale, in **Figure S9** we depict three cases based on *in silico* data where each circle was incorrectly detected (FP) and its corresponding true positive circle. The circle A is a case of exact duplication, and thus both have the exact value of ΔCJ=0 and *p*_adj_=1. On the other hand, circles B and C stem from a partial duplication, where one extreme of the circle does not share coordinates with the simulated circle, and thus the FP circles show higher ΔCJ values and lower configuration probabilities (**Table 2**).

**Table 2.** ΔCJ and *p*_adj_ ΔCJ values of 3 circle detection examples. For each circle, the number of assigned reads, ΔCJ and *p*_adj_ ΔCJ are calculated for the true positive and FP circles. TP: true positive; FP: false positive.

| Circle | Condition | Reads (L\|R) | $\Delta CJ$ | $p_{adj}$ $\Delta CJ$ |
| --- | --- | --- | --- | --- |
| chr6:39,754,059-39,754,823 | TP | 22 \| 22 | 0 | 1 |
| chr6:39,754,063-39,754,823 | FP | 22 \| 22 | 0 | 1 |
| chr3:140,171,091-140,172,928 | TP | 28 \| 27 | 0.018 | 1 |
| chr3:140,171,091-140,174,822 | FP | 28 \| 0 | 1 | $5.7 \cdot 10^{-8}$ |
| chr17:6,795,106-6,797,132 | TP | 25 \| 29 | 0.074 | 1 |
| chr17:6,794,297-6,797,033 | FP | 7 \| 39 | 0.69 | $1.1 \cdot 10^{-5}$ |

Assuming that the number of reads assigned follows a binomial distribution *B*(*n, p* = 0.5), we can set a circle with a high imbalance (0 or 1 reads assigned on one side) as an incorrect circle with *α* = 0.05 if it contains 9 or more reads (described in Materials and Methods). Additionally, for each circle with *k*_*L*_ reads assigned to the left, and *k*_*R*_ reads assigned to the right, we can compute the probability *p* of this configuration, assuming a binomial *B*(*n* = *k*_*L*_ + *k*_*R*_, *p* = 0.5) distribution. Based on this, we will compute a series of metrics: (1) proportion of circles with *≥* 9 reads, (2) mean and median ΔCJ and (3) ratio of circles with *p*_adj_ *<* 0.05, where probabilities are adjusted by the Benjamini-Hoechberg method.

To ensure that the metric is a correct proxy for the accuracy of circle detection, we applied it to *in silico* data. To that end, in **Figures S10A** and **B** we show the ΔCJ distribution of TP reads and FP reads for eccDNA and circRNA, respectively. Overall, simulated circles follow a right-tailed distribution of ΔCJ, with median values smaller than 0.1 for both eccDNA and circRNA, showing that most circles have a balanced left/right read distribution. The ΔCJ distributions arising after filtering showed no statistically significant differences, neither for eccDNA (Kruskal-Wallis test, H=2.2854, *p*_adj_=0.683) nor for circRNA (H=0.0229, *p*_adj_=0.999)

In eccDNA, the unfiltered FP circles showed a wider ΔCJ distribution, with a mean value of 0.38, and many of them with ΔCJ=1 (75th percentile is 1). Similarly, circles that still remain FP after filtering showed a high median ΔCJ, even higher than the unfiltered circles. To better understand individual the filtering effect, circles remaining FP after *filter-split* showed a lower ΔCJ than circles remaining after *filter-duplicates*. This may indicate that *filter-split* removes high-ΔCJ circles, whereas *filter-duplicates* removes low-ΔCJ circles. This effect is also observable based on their assignment probability: there is a higher proportion of circles retained after *filter-duplicates* that have *p*_adj_ < 0.05 (77%), indicating that most of these are unbalanced, in contrast to the proportion of circles retained after *filter-split* (19%) (**Supplementary Material 19**). Thus, most of these low-ΔCJ circles are, expectedly, duplications of simulated circles, and thus may contain a balanced left/right-assigned reads ratio. On the other hand, high-ΔCJ circles may represent new circles, which escape duplicate-filtering, and which that contain an imbalanced left-right read ratio, as explained before.

Interestingly, the results on circRNA are different. All circles retained as FP after filtering remain with a low ΔCJ; with the exception of *filter*, which has a similar ΔCJ distribution as in eccDNA. These distributions are likely because most of these circles may be assigned uniquely by segemehl, which contains a extremely high amount of duplicated circles, most of which may be even remain after *filter-duplicates*. Therefore, the remaining circles contain a low ΔCJ. Interestingly, the ratio of FP circles with *p*_adj_ < 0.05 is higher after filter-split (37%) than after *filter-duplicates* (8%) indicating that many of the removed circles by *filter-duplicates* probably have a biased left/right read ratio (**Supplementary Material 20**).

#### Circle capture method greatly affected detection quality

After establishing ΔCJ as a proxy measure to evaluate the fit of the circular detection, we wanted to observe the variations in ΔCJ for the different circle capture methods in biological data. **Figure 5** shows ΔCJ values for all 4 circle capture methods in each circle detection tool and the four filtering methods.

**Figure 5.**
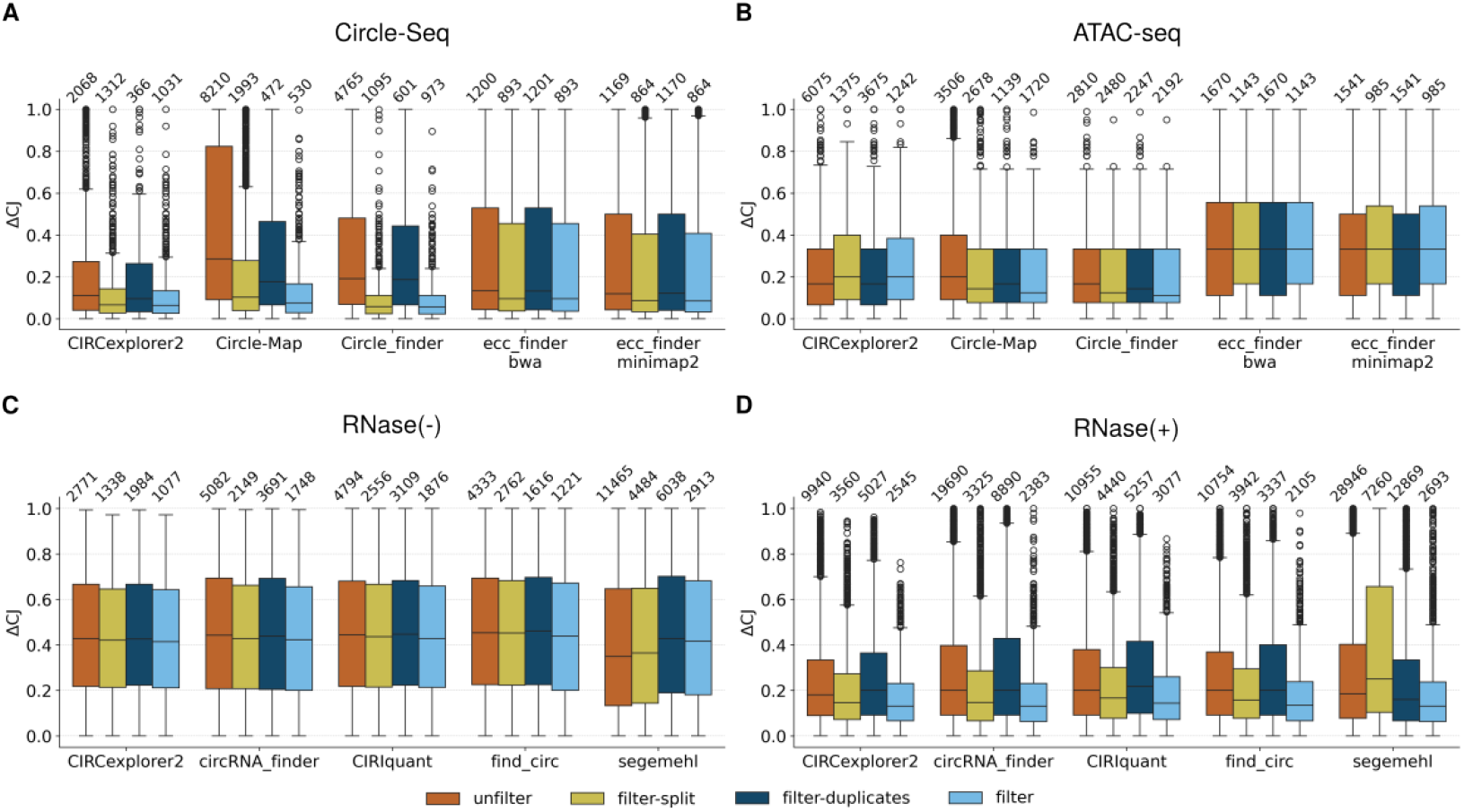
Performance analysis of detection software for eccDNA and circRNA identification in biological datasets. Boxplots of ΔCJ values for (**A**) Circle-Seq and (**B**) ATAC-seq data for eccDNA, and (**C**) RNase(−) and (**D**) RNase(+) data for circRNA, under four filtering conditions: *unfilter, filter-split, filter-duplicates*, and *filter*.

Regarding eccDNA detection tools, similarities between individual tools were visible again. For instance, ecc_finder-bwa and ecc_finder-minimap2 showed a similar detection pattern based on the ΔCJ for both Circle-Seq and ATAC-seq. In Circle-Seq, the filtering effect on the ΔCJ is more apparent than for ATAC-seq. Generally, there are more circles and with lower ΔCJ values remaining after *filter-split*, compared to *filter-duplicates*. ΔCJ values for *filter* are similar, if not lower, to *filter-split*, suggesting again that these two filtering strategies are optimal also for biological datasets.

Additionally, ATAC-seq has a tendency to capture more circles than Circle-Seq, as mentioned before. Interestingly, in tools and filter combinations where the circle number is similar or higher, ΔCJ are higher in ATAC-seq; with few exceptions like *filter-duplicates* in Circle_finder and CIRCexplorer2. Also, ΔCJ for ecc_finder tools are markedly higher in ATAC-seq compared to Circle-Seq. Therefore, it is likely, as mentioned in the previous section, that the higher number of circles captured by ATAC-seq may not correlate with circle quality.

Focusing on circRNA, there is a clear difference in the distribution of ΔCJs: RNase(−) has a stable ΔCJ at around 0.4 (differences in *filter*, Kruskal-Wallis test: H = 3.49, *p*=0.48); while effects across filters are more pronounced in RNase(+) (differences in *filter*, Kruskal-Wallis test: H = 17.72, *p*=0.0013), which shows a clearly reduced ΔCJ compared to RNAse(−). In this case too, compared with eccDNA, a lower number of detected circles correlates with the ΔCJ value (e.g. with different filters) indicating that the two-filtering process tends to retain circles with lower ΔCJ. Additionally, and similarly to eccDNA, circles remaining after *filter-split* show lower ΔCJs than after *filter-duplicates*. Therefore, these results suggest that RNase treatment may enhance detection accuracy (**Supplementary Material 21, Supplementary Material 22**).

#### Rosette tool combination yielded the best trade-off between detection accuracy and number of circles

The final aspect of this analysis is centered on evaluating tool combinations. Firstly, we assessed how ΔCJ varies depending on the combination of tools. To this end, **Figure S11** shows the ΔCJ values for each combination (**Supplementary Material 23**), while **Figure 6** illustrates the difference in ΔCJ between the *Rosette* combination and all other combinations (ΔΔCJ).

**Figure 6.**
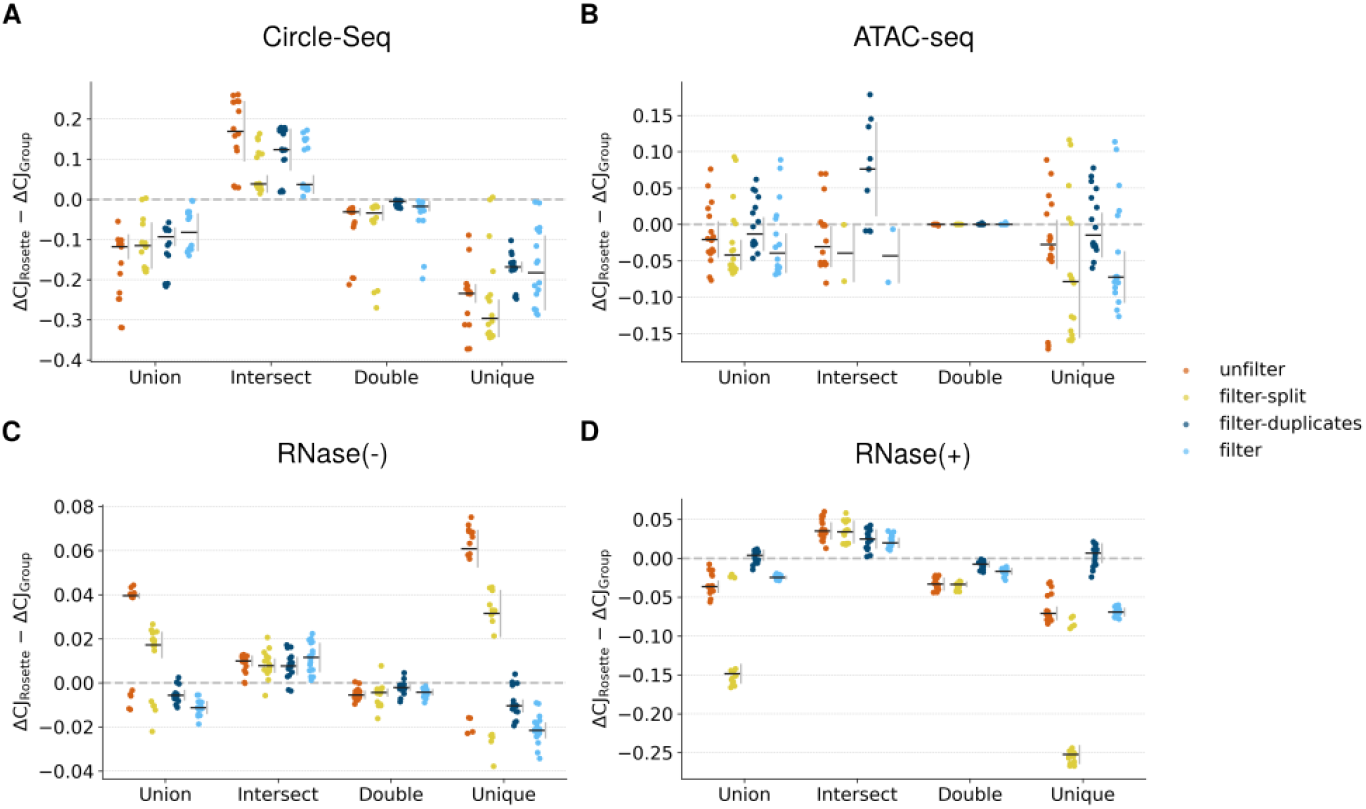
Performance analysis of software combinations for eccDNA and circRNA identification in biological data. Strip plot of ΔΔCJ values (ΔCJ_Rosette_ - ΔCJ_Group_) for different tool combinations—*Union, Rosette, Intersect, Double*, and *Unique*—evaluated on biological data, with comparisons made against the *Rosette* combination. Results are shown for (**A**) Circle-Seq and (**B**) ATAC-seq data for eccDNA, and (**C**) RNase(−) and (**D**) RNase(+) data for circRNA, under four filtering conditions: *unfilter, filter-split, filter-duplicates*, and *filter*.

From this analysis, although results vary considerably based on the filter and circle enrichment method, we observed a general trend: the rest of tool combination strategies consistently yielded ΔΔCJ < 0, meaning that the *Rosette* combination typically has a lower ΔCJ compared to other combinations. This trend is particularly evident when comparing *Rosette* with *Unique*, where ΔΔCJ is the most pronounced. Regarding filtering, as previously discussed, the lowest ΔCJ values are generally achieved with the *filter* strategy, followed by *filter-split*.

The only exception exhibiting ΔΔCJ > 0 occurred with the *Intersect* combination. This result is expected, as *Intersect* exclusively selects circles detected consistently by all tools. However, *Intersect* detected significantly fewer circles, about half compared to *Rosette* (**Figure S12**), suggesting that the *Rosette* combination might provide a better balance between the number of detected circles and detection quality.

The ΔCJ values observed between *Rosette* and *Double* combinations were quite similar, with *Rosette* generally having slightly lower ΔCJ values. Nevertheless, this relationship depends on the applied filter (e.g., in RNase(+) ΔΔCJ varies from −0.005 to −0.04). This similarity in ΔCJ between *Double* and *Rosette* is to some extent surprising, considering that *Double* excludes *Intersect* circles. This showcases that the weight of *Rosette* ΔCJ comes from double circles instead of *Intersect* ones.

Finally, we investigated how the number of tools affects ΔCJ. In **Figure S13** ΔCJ is plotted against the number of tools in each tool combination, circle enrichment technique, and filter. Generally, although particularly for *Rosette*, a higher number of tools slightly increased ΔCJ. In certain cases (especially for *Double*), combinations involving more tools produce ΔCJ values midway between individual combinations.

Our interpretation of this phenomenon is that adding more tools could introduce biases from tools such as segemehl or ecc_finder, which, as previously observed, may have lower detection accuracy. Thus ΔCJ values are higher, thereby slightly increasing the combined circles’ ΔCJ. However, this increase in ΔCJ from including additional tools is considerably smaller than the effect introduced by the filtering method or circle enrichment technique.

We observed the opposite effect with *Intersect*, where the addition of more tools skewed the ΔCJ distribution towards lower values. As previously explained, this occurs because *Intersect* retains only circles consistently detected across multiple tools, inherently selecting those with greater detection reliability and thus lower ΔCJ.

## Discussion

The accurate identification of eccDNA and circRNA is essential for understanding their biogenesis, functions, and implications in diseases. These circular molecules play significant roles in gene regulation and are increasingly recognized for their involvement in various pathological conditions, including cancer and neurodegenerative disorders, highlighting their potential use in disease diagnosis and monitoring (Turner et al., 2017; Zhang et al., 2018; Iparraguirre et al., 2017; Noer et al., 2022). However, the lack of standardized evaluation frameworks and consensus protocols for their detection poses a significant challenge, often leading to inconsistencies across studies. To address this issue, we conducted a systematic benchmarking of existing detection methods using both *in silico* and biological datasets, aiming to clarify their strengths, limitations, and practical applicability.

Detection performance varied significantly depending on the type of circular molecule analyzed. Regarding eccDNA, we observed that ecc_finder-minimap2 and CIRCexplorer showed robust results, followed by Circle-Map and Circle_finder. However, applying any filtering strategy led to similar outcomes across all tools. These observations are consistent with findings reported by Li et al. (2024) and Gao et al. (2024), who also identified Circle-Map and Circle_finder as top-performing tools. Furthermore, Gao et al. (2024) reported poor performance of ecc_finder-bwa, aligning with our results and indicating partial reproducibility between studies.

In the case of circRNAs, the comprehensive analysis by Zeng et al. (2017), although not entirely replicable in our study, suggests a correlation between the number of circles detected and the F1-score for each tool. Conversely, Vromman et al. (2023) observed a clear detection bias, finding segemehl among the tools with lower detection rates, contrasting sharply with our results. This discrepancy could arise from several factors, including the fact that segemehl, initially designed as an aligner, was later integrated into circRNA analysis workflows; consequently, its performance might be highly dataset-dependent. Additionally, Liu et al. (2022) analyzed the effect of coverage similar to our study, although they did not observe significant impacts of coverage on detection, possibly due to their analysis not including higher coverage levels like ours, where we indeed observed an increase in FPs. They also reported better performance for CIRI and circRNA_finder compared to our findings, potentially explained by their use of CIRI-simulator (Gao et al., 2015) for circle simulation, which may inherently bias results in favor of these tools.

Focusing on biological datasets, we found the library preparation method to be one of the most influential factors for both eccDNA and circRNA detection—specifically, ATAC-seq versus Circle-seq for eccDNA, and RNase+ versus RNase− for circRNA. For eccDNA, we detected a greater number of circles using ATAC-seq compared to Circle-seq, conflicting with results reported by Gao et al. (2024), who observed significantly higher detection rates (up to two orders of magnitude) for Circle-seq. This discrepancy might be attributed to sample origin or other internal methodological factors, though the exact reason remains unclear.

In contrast, our circRNA findings align well with studies such as Zeng et al. (2017), who reported increased circle detection following RNase treatment (approximately 1.5 to 3 times higher). Nonetheless, we observed that circle detection numbers strongly depend on the tools employed. One potential confounding factor influencing this variability is the alignment algorithm, which differs among tools. To minimize this variability, we standardized the aligners used as much as possible—employing STAR for circRNA and bwa for eccDNA detection. However, disparities in results still indicated significant aligner-specific factors influencing circle detection accuracy.

We also conclude that targeted amplification methods notably affect detection outcomes, potentially introducing additional biases. For example, Rolling Circle Amplification (RCA), used in Circle-seq, can lead to either overrepresentation or underrepresentation of circles (Yu et al., 2024). Moreover, such amplification methods may impede the analysis of epigenetic signals, which are linked to gene expression variations and regulatory alterations. Consequently, non-targeted sequencing approaches such as ATAC-seq and RNA-seq are gaining interest, although further investigation is necessary to fully understand and address potential biases inherent in these methods.

Delving deeper into the analysis, several structural effects deserve discussion, particularly the issue of FPs. Initially, we observed that coverage substantially influences FP detection in *in silico* data, a result whose applicability to biological data remains unclear but warrants consideration in analyses.

We aimed to understand the origin of these FPs. Zeng et al. (2017) defined FPs as circles detected in RNase(−) treatment but not in RNase(+), which theoretically amplifies the circular RNA signal. Our findings highlight the need for further exploration of FP origins, particularly in *in silico* datasets, to enhance their applicability in biological studies. We propose that FPs may originate from two main sources: computational and biological effects. Computational effects include coverage-related issues, where increased sequence detection near the CJ may occasionally lead to incorrect identification due to partial flanking or sequencing errors, creating artificial circles. Additional computational factors encompass sequence length, the aligner used, and the detection algorithm itself. Biological effects primarily involve amplification methods—techniques excluding linear DNA or RNA can reduce FPs by preventing amplification of sequences resembling CJ, which may result from alternative splicing or genomic mutations and rearrangements that do not yield genuine circles. Consequently, methods involving purification steps (such as Circle-Seq for eccDNA and RNase(+) for circRNA) generally provide superior results compared to those lacking this step.

Another crucial factor affecting the analysis and significantly reducing FPs is the appropriate use of filters. Both *in silico* and biological datasets demonstrated a considerable decrease in detected circles upon filtering. Our findings align with Vromman et al. (2023), who reported substantial reduction in circles with *≥*5 reads mapped to the BSJ.

In our analysis, apart from applying a filter based on counts (*≥*2 reads), we also implemented a duplicate filter. Generally, both filters performed adequately, though the filter-split occasionally yielded slightly superior results. A potential reason is that many tools do not provide precise CJ coordinates; rather, they often detect circles at adjacent coordinates, typically supported by low read counts (1 or 2) and occasionally mismatches that cause partial misalignment. Thus, many spurious circles escape duplicate filtering, making filter-split more effective in such cases.

An additional intriguing observation from our *in silico* analysis pertains to circle length. We noted a marked decline in detected circles around 340 bp in both eccDNA and circRNA, possibly corresponding to dinucleosome length. However, since *in silico* data are independent of biological factors, this finding is surprising. It could be attributed to detection biases such as those observed in ecc_finder, which fails to detect shorter eccDNAs (<400 bp), an issue also reported by Li et al. (2024).

Lastly, repetitive sequences significantly impact circle detection. Gao et al. (2024) observed pronounced effects on detecting reads associated with LTR, SINE, and LINE elements, depending on whether the sequencing technology involved long or short reads, albeit representing a small fraction of total reads (0.1% for short reads). Our *in silico* results showed F-scores slightly above 0.8 for eccDNA and between 0.8 and 1 for circRNA for LINEs and SINEs, suggesting a slightly higher yet potentially problematic detection fraction in biological datasets due to the repetitive nature of these sequences. The most pronounced effect occurred with satellite sequences, where detection accuracy declined sharply. This phenomenon may substantially impact eccDNA detection in centromeric regions, which, due to their repetitive nature, could be inaccurately captured, potentially leading to significant functional loss in analyses.

A significant limitation of current benchmarking studies is the absence of robust metrics for evaluating accurate circle detection in biological data. Often, comparisons rely on the number of detected circles or similarity between tools, neither of which provide true validation. Alternative methods such as external validations using qPCR or similar techniques cannot scale effectively beyond a few circles.

In this study, we employed the ΔCJ metric, developed under the assumption that reads assigned to the CJ will equally distribute between its left and right sides. Consequently, significant deviations from this symmetry indicate potential incorrect assignments. After confirming the rationale through specific examples and verifying its applicability in *in silico* data, we found that the use of ΔCJ in biological datasets provides valuable insights for evaluating tool effectiveness, purification methods, and filtering strategies.

However, it is crucial to recognize the conceptual and practical limitations of ΔCJ. For instance, some FP circles might escape detection due to factors unrelated to symmetry imbalances, limiting the metric to detecting only certain types of FPs. Additionally, the reliability of this metric requires a relatively high number of reads (preferably more than 20-30), restricting its effectiveness for detecting circles with low read counts. Nevertheless, one advantage is that high-read FPs with significant asymmetry may potentially be reassigned to circles sharing one CJ coordinate. Lastly, we observed that for circRNAs, the metric is noisier, possibly because many circRNAs have boundaries defined precisely at intron-exon junctions, conflicting with the expected randomness of read distribution. To improve on this limitations a more robust theoretical framework of circle read mapping to CJ that accounts for sequencing depth and circle length may be necessary.

One advantage of incorporating multiple detection tools in analyses is the improvement in eccDNA and circRNA detection, especially considering analysis workflows like those implemented in nf-core, which facilitate integration of multiple tools (Ewels et al., 2020). This observation aligns with Hansen (2018), who identified commonly detected circles as *bona fide* circles. Other studies have similarly highlighted the variability in detection consistency among tools. For instance, Vromman et al. (2023) reported that nearly 50% of circles detected were unique to a single tool—though the exact percentages varied considerably by tool—while approximately 10% were consistently detected across ten or more tools. Likewise, Li et al. (2024) suggest that including two or more tools can enhance the robustness of circle detection.

Vromman et al. (2023) validated circles consistent across multiple tools, observing that employing two or more tools notably reduced FPs. However, while this outcome could theoretically extend to eccDNA, direct validation for eccDNA is lacking, and additional validation measures for various tool combinations are needed. Our benchmark results underscore that the *Rosette* combination strategy achieves the best balance between the number of circles reported and FP reduction.

The rationale for the *Rosette* strategy is that circles detected by multiple tools are likely to represent true positives, whereas those identified by a single tool should be interpreted cautiously. Among all tested combinations, *Rosette* exhibited superior performance in simulated data by significantly lowering the FP rate while maintaining high sensitivity, even without additional filtering. Similarly, in biological data, it enhanced detection capacity without compromising accuracy. Nevertheless, further validation using deeper analysis of reads associated with detected circles would bolster the reliability of this approach.

Additionally, we observed that the effectiveness of combined detection strategies depends on the individual accuracy of each tool. This aligns with Vromman et al. (2023), who found that the FP rate for combined tools approximates an average of the individual FP rates. Thus, the choice of combination strategy (particularly between *Rosette* and *Intersect*) depends on the analytical goal. In clinical settings or environments requiring higher positive predictive value, *Intersect* may be more advantageous despite detecting fewer circles. In contrast, *Rosette* may offer a sufficiently robust and reliable circle set suitable for broader research applications.

## Methods

### eccDNA and circRNA detection software

In this study, we compared 5 eccDNA detection software–Circle-Map (v1.1.4), CIRCexplorer2 (v2.3.8), Circle_finder, ecc_finder-bwa (v1.0.0), and ecc_finder-minimap2 (v1.0.0)–and 5 circRNA detection software–CIRCexplorer2 (v2.3.8), circRNA_finder (v1.2), CIRIquant (v2.1.0), find_circ (v1.2), and segemehl (v0.3.4). All of the software tools used in this study are integrated into the nf-core framework, with the exception of ecc_finder. We performed the detection of eccDNA using nf-core/circdna (v1.1.0) (Schreyer et al., 2024), and the detection of circRNA using nf-core/circrna (dev) (Digby et al., 2023). Sequence reads were aligned to the human reference genome GRCh38 (NCBI) and the mouse reference genome GRCm38 (Ensembl).

For read alignment, eccDNA sequencing reads were mapped to the reference genome using BWA (v0.7.17-r1188), while circRNA sequencing reads were aligned using STAR (v2.7.11b). Alignments were performed against the human reference genome GRCh38 (NCBI) and the mouse reference genome GRCm38 (UCSC)

Although all software’s detection is base on identifying the circular junction, their strategies for identifying circles is different.

#### eccDNA

CIRCexplorer2 is the upgraded version of CIRCexplorer (Zhang et al., 2014). It was primarily developed to detect circRNA, but it can also identify eccDNA. CIRCexplorer2 integrates additional aligner options beyond the original TopHat2 (Kim et al., 2013), including STAR (Dobin et al., 2013), MapSplice (Jeck et al., 2013) and segemehl, to accommodate different RNA-seq mapping preferences. CIRCexplorer2 alignes reads to the reference genome using various aligners and detects non-colinear reads are detected. Next, CIRCexplorer2 analyzes these non-colinear alignments to detect the exact location of the CJ. Additionally, CIRCexplorer2 reanalyzes reads that were originally mapped to linear exon-exon junctions. For circRNA, it also performs de novo assembly of linear reads to discover novel exons and splicing events. Additionally, unmapped reads are realigned to capture any missed circular structures. In this study, DNA was mapped using BWA (Li and Durbin, 2009), and RNA was mapped using STAR. For *in silico* circRNA detection, we selected an intermediate non-annotated file due to the lack of detected circRNAs in the annotated output generated by the nf-core CIRCexplorer module.

Circle-Map identifies eccDNA breakpoints by utilizing discordantly mapped reads and mapping soft-clips using probabilistic models. First, Circle-Map detects eccDNA candidate reads, including discordant read pairs, soft-clipped reads, and hard-clipped reads, using the BWA aligner. Next, it constructs a breakpoint graph based on these candidate reads. Finally, soft-clipped reads are realigned using a probabilistic model to accurately determine the eccDNA breakpoints and achieve nucleotide-level resolution.

Circle_finder is designed to identify eccDNA from whole-genome sequencing (WGS) and ATAC-seq data by analyzing read pairs using BWA. It collects all read pairs where one read maps uniquely to the genome in a contiguous manner and the other read maps as a split read flanking the mapped read. The start of the split read and the end of the contiguous read are then annotated as the start and end points of the eccDNA

ecc_finder detects circular breakpoints based on discordant reads and split reads. Once discordant reads and split reads are identified, only reads spanning the same boundary are retained to define the breakpoint. By default, ecc_finder uses the BWA aligner, but Minimap2 (Li, 2018) can also be applied for alignment.

#### circRNA

circRNA_finder uses STAR to directly identify chimeric junction reads from RNA-Seq data. After the initial alignment, the algorithm filters these chimeric reads to detect potential CJs based on predefined criteria. The filtering process includes evaluating the uniqueness of the mapped reads, allowing for a limited number of mismatches, and ensuring that the distance between splice donor and acceptor sites is within a specified range.

CIRIquant extends the functionality of CIRI2 by implementing a pseudo-reference-based approach for circRNA detection. Initially, reads are aligned to the reference genome using BWA, and unmapped reads are considered as potential circRNA candidates. CIRCquant then uses CIRI2 to detect circRNAs, but it also supports BSJ bed files created by other software. A pseudo-reference consisting of circular sequences is generated, and all circular candidate reads are aligned to the pseudo-reference using HISAT2. Reads that map concordantly within a 10 bp region of the BSJ are classified as circular reads. Additionally, CIRIquant can perform RNase correction, linear RNA quantification, and circRNA differential expression analysis.

find_circ uses a segment-based approach to identify circRNA. First, reads are mapped to the reference genome using Bowtie2, and reads that align contiguously are discarded. From the remaining reads, 20 nucleotides from both ends are extracted and aligned to obtain unique anchor positions within spliced exons. Anchors that align in the reverse orientation are identified as circRNAs. The anchor alignments are then extended to the BSJ, flanked by GU/AG splice sites.

Segemehl is able to identify multiple types of splice junctions. It aligns RNA-Seq reads to a reference genome while accounting for complex splicing patterns that are characteristic of circular RNAs. The software detects chimeric reads that span back-splice junctions, where the 3’ end of one exon is joined to the 5’ end of another in a circular fashion.

### *in silico* datasets

#### CircleSim. CircleSim

is a specific simulation software for circular and linear reads. It is implemented in Python 3 and consists of three modules: 1) coordinates: generates coordinates based on a provided length distribution; 2) reads: simulates circular or linear short-read sequencing; and 3) join: merges circular and linear FASTQ files.

The coordinates module selects a chromosome at random based on length-associated probabilities. For both DNA and RNA, the first nucleotide of the CJ is randomly chosen across the entire genome and transcriptome, respectively. The position of the second nucleotide is determined by the length of the circular region, which can be modeled using either a uniform or a lognormal distribution. The uniform distribution is defined by specified minimum and maximum lengths, while the lognormal distribution is characterized by its mean and standard deviation, with options to set minimum and maximum lengths as well.

The reads module simulates sequencing based on the read and insert lengths, and a coverage defined as:

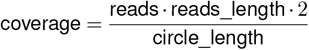

For circular molecules, a nucleotide within the circle is selected randomly, and the distance to the CJ determines whether the read is concordant, discordant, or a split-read. Concordant reads are those mapped in the expected orientation, discordant reads are mapped in the opposite orientation, and split-reads have an unmapped portion because they span across the CJ. CircleSim includes an option to force reads be near the CJ. For linear molecules, the start of the read is randomly selected between the start of the sequence and the position defined as the end of the sequence minus the insert length, ensuring that all reads are concordant.

The source code is released under the MIT license and is freely available at https://github.com/ZabalaAitor/CircleSim.

#### in silico datasets

We used *CircleSim* to generate 1,000 eccDNAs and circRNAs from the canonical chromosomes and transcripts for the GRCh38 version (NCBI) of the human genome, respectively. The circles were simulated with a size distribution ranging from 175 to 10,000 bp in a log-normal distribution, with a mean of 1,000 and a standard deviation of 1. The reads were simulated based on short-read sequencing with a read length of 150 bp, an insert length of 500 bp, a sequencing error rate of 0.001, and a mutation rate of 0.01 under the Kimura mutation model, with coverage depths of ×5, ×7, ×10, ×15, ×20, ×30, ×50, ×70, and ×100.

### Biological datasets

We downloaded a dataset (SRR6315430) from human muscle tissue, where eccDNA was enriched and sequenced using Circle-Seq (Møller et al., 2018). We also used a dataset (EGAS00001005873) generated by The Chinese University of Hong Kong (CUHK) Circulating Nucleic Acids Research Group from a knockout mouse models with deficiencies in deoxyribonuclease 1 like 3 (DNASE1L3), where cell-free eccDNA (cf-eccDNA) was sequenced using ATAC-seq (Sin et al., 2022) sequencing. For circRNA, we downloaded two HELA datasets (Gao et al., 2015): an original sample (SRR1637090) and the corresponding sample after RNase R treatment (SRR1636986). We used ‘prefetch’ and ‘fasterq-dump’ from the SRA Toolkit (https://github.com/ncbi/sra-toolsv2.9.1) to download and convert the SRA files to FASTQ format.

The number of reads associated to each dataset is the following: Circle-Seq - 12,829,402; ATAC-seq - 12,287,079; RNase(−) - 35,685,310; RNase(+) - 23,505,713.

### Circle filtering strategies

To improve the reliability of predicted circular elements, we implemented four filtering strategies: *unfilter, filter-split, filter-duplicates, filter*. These strategies aim to reduce FPs and emphasize consistently detected circles across different algorithms. The filtering strategies used in this study are described below:

- ***unfilter*** : Includes all raw detections without applying any filtering criteria.
- ***filter-split*** : Retains only circles supported by at least two split reads.
- ***filter-duplicates***: Removes overlapping circles, where two circles are considered overlapping if their coordinate-defined regions share at least one base. Among overlapping circles, the one with the highest number of supporting split reads is retained. If split read information is unavailable, the longest circle is kept.
- ***filter*** : Applies both split-read filtering and duplicate removal. Specifically, it first retains circles with at least two supporting split reads, and then removes overlapping circles as described above.

All tools used in this study provide split read information, except for *CIRCexplorer2* in the context of *in silico* circRNA detection, in which split information is not considered.

### Circle combination strategies

To assess the reliability of circle detection across multiple tools, we evaluated the performance of different tool combination strategies (**Figure 7)**:

**Figure 7.**
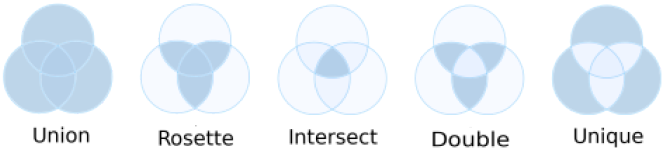
Description of the combining strategies. Visualization of the different combining strategies: *Union, Rosette, Intersect, Double*, and *Unique*.

- ***Rosette***: Includes circles detected by at least two different tools in a combination of three or more tools.
- ***Union***: Includes all circles detected by any of the tools.
- ***Intersect*** : Includes only circles detected by all tools in the set.
- ***Unique***: Includes circles detected by only one tool.
- ***Double***: Includes circles detected by at least two tools, excluding both *Unique* and *Intersect* detections.

The combinations shown in the main plots are based on sets of three or more tools, in accordance with the minimum requirement of three tools for the *Rosette* strategy. However, two tool combinations for *Union, Intersect*, and *Unique* are included in the Supplementary Material for completeness.

### *in silico* statistical analysis

#### Base metrics

A threshold of 20 bp between the analyzed circles was applied to determine if two circles were considered the same. Detection accuracy in simulated data was analyzed using precision, recall, and F-score metrics.

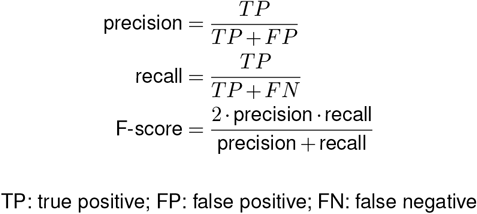

TP: true positive; FP: false positive; FN: false negative

#### Circular junction detection precision

To evaluate the offset of the CJ detection, we calculated the ratio of number of circles detected with an offset of 1 bp (eccDNA_T=1_ for eccDNA and circRNA_T=1_ for circRNA) and the maximum number of circles detected (eccDNA_max_ for eccDNA and circRNA_max_ for circRNA). Higher ratios indicate that most circles are detected with only a small offset of 1bp, while lower ratios indicate that a higher offset is required.

#### Length distribution

The distribution of eccDNA and circRNA lengths was analyzed and compared using the Kolmogorov-Smirnov (KS) test. To account for the non-uniformity of the distributions, we first examined the full range of circle lengths (175–10,000 bp). We then focused specifically on short circles, defined as those within the 175–1,000 bp range. Finally, we identified a specific length interval—referred to as the “square” range—where the observed distribution was lower compared to the simulated distribution. The circular length distribution was plotted using both absolute and relative counts. The relative counts were calculated by normalizing against the counts from the simulated data. To enhance the clarity of the distribution plots, a sliding window of size 5 was applied to smooth the distribution curves. This technique allowed for more precise visualization of the trends across different tools, facilitating the comparison between predicted lengths and the ground truth, represented by the simulated circles. For this analysis, we used *in silico* datasets generated at ×30 coverage, ensuring consistency across comparisons.

#### Repeat element annotation

For the repeat element analysis, we utilized the GRCh38 (NCBI) genome assembly in conjunction with RepeatMasker open-4.0.5 and the Repeat Library (January 31, 2014). The RepeatMasker software, using the specified library, allowed us to identify and classify repetitive elements within the genome. Repeat elements were categorized into several classes, including LINEs (Long Interspersed Nuclear Elements), *SINEs* (Short Interspersed Nuclear Elements), *DNA* (DNA transposons), *satellite* (satellite DNA), and *other* elements such as *Long Terminal Repeats (LTR), simple repeats, low-complexity regions, snRNA*, and *unknown* elements. Junctions that did not overlap with any annotated repeat were labeled as *non-repetitive* (Ø). For this analysis, we used in silico datasets generated at ×30 coverage, ensuring consistency across comparisons. This analysis was performed to ensure comprehensive annotation of repeat sequences at each coordinate of the CJ. For this analysis, we used *in silico* datasets generated at ×30 coverage, ensuring consistency across comparisons.

#### Genomic element annotation

For the genomic element analysis, we used the GFF annotation file corresponding to the GRCh38 version (NCBI) of the human genome. This analysis aimed to provide comprehensive annotation of genomic elements at each CJ coordinate. Initially, all coordinates were classified as *intergenic*. Subsequently, coordinates were annotated based on detailed genomic features, including *3’-UTR, 5’-UTR*, and *other* elements such as *start codon, stop codon*, and *selenocysteine positions*. Coordinates not annotated in this step were then evaluated for overlap with *exon* regions. Those that still remained unannotated were next checked against *intronic* regions. Finally, any coordinates not matching any of the above categories were retained as *intergenic*. For this analysis, we used *in silico* datasets generated at ×30 coverage, ensuring consistency across comparisons.

### Biological statistical analysis

#### Similarity analysis

eccDNA and circRNA detection results were visualized using UpSet plots (Lex et al., 2014), which allow visualization of overlapping circles defined with a threshold of 20 bp.

#### Circular junction difference (ΔCJ)

When a read spans the CJ, the split, unmapped portion of the read can be associated either with the left or right side of the junction. Assuming equal probability for reads to associate with either side of the CJ, we hypothesized that circles showing disproportionate read association on one side are likely FPs, with these reads potentially originating from other circles.

Based on this hypothesis, if *N* is the number of reads spanning a circle’s CJ, *k*_*L*_ is the number assigned to the left side, and *k*_*R*_ = *N* −*k*_*L*_ is assigned to the right, we defined the metric ΔCJ as:

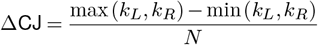

To obtain *k*_*L*_ and *k*_*R*_, we extracted reads overlapping windows centered on the circle start and end positions, respectively. Each window extended *N*_offset_ = 20 nucleotides upstream and downstream of the junction coordinate to capture all potentially spanning reads. Unique read identifiers were collected within these windows from the aligned BAM file using the pysam package for efficient read retrieval. Counts were then derived for each side. The total number of CJ reads, *N*, corresponds to the union of reads found in both windows.

Values of ΔCJ close to 0 indicate a balanced read distribution, whereas ΔCJ = 1 means that all reads associate with only one side of the junction.

While informative, this metric has limitations, particularly at low *N*, where extreme ΔCJ values are common. To address this, we modeled the probability of observing a particular distribution of reads (*k*_*L*_ and *k*_*R*_) to identify circles where such probabilities are exceedingly low.

The number of reads associated with one side of the CJ, *X*, can be modeled using a binomial distribution: *X* ~ *B*(*N*, 0.5). Thus:

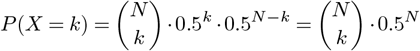

Considering that assignment to either side of the CJ is arbitrary, the final distribution must account for symmetry:

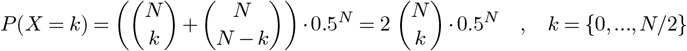

Thus, the cumulative probability of assigning up to *k* reads out of *N* to one side of the CJ is:

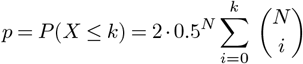

For even *N* values, the cumulative distribution can exceed 1 due to double-counting at the midpoint *P* (*X* = *N/*2) For example, when *N* = 6 and *k* = 3, the cumulative probability is: 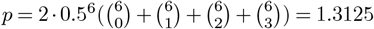. This happens because *P* (*X* = 3) is counted twice. In contrast, when *N* = 7, for *k* = 3, 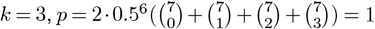 as the complement of 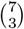 is 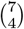. Thus, in even cases where *k* = *N/*2, the cumulative probability is set to 1 since midpoints are not relevant for identifying outliers, which are our primary interest.

Under these conditions, the minimum required to achieve a cumulative probability *P* (*X* ≤ 1) below 0.05 is: 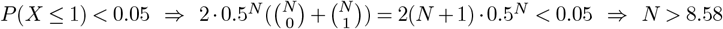

The solution was retrieved numerically. For *P* (*X* ≤ 1) *<* 0.01 the solution is *N >* 11.25.

Using these findings, we defined three metrics for evaluating circle detection quality: (1) the proportion of circles with *N* ≥ 9, (2) ΔCJ for selected circles, and (3) the proportion of circles fulfilling *p <* 0.05 based on probabilities derived from observed *k* and *N* values. For the third metric, we applied the Benjamini-Hochberg correction, designating circles with adjusted probabilities (*p*_adj_ *<* 0.05) as significantly skewed compared to the expected baseline distribution.

## Computational resources

All circular detection analyses were conducted using on a single computing node with 256 GB of RAM and 24 cores.

## Data availability

All the data generated is available at 10.5281/zenodo.15783793.

## Code availability

All the code is available at 10.5281/zenodo.15783795 and https://github.com/ZabalaAitor/benchmarking

### Author contributions

Author contributions are detailed according to the CRediT taxonomy. AZ, AMA, and DO contributed to the conceptualization of the study. Methodology was developed by AZ, AMA, IPL, and DO. Software development and implementation were carried out by AZ and AMA. Validation and formal analysis were performed by AZ and AMA. AZ was responsible for the investigation. Resources were provided by AZ, AMA, and DO. Data curation was handled by AZ and AMA. All authors—AZ, AMA, IPL, and DO—contributed to writing the original draft, as well as to reviewing and editing the manuscript. Visualization was carried out by AZ. Supervision was provided by AMA, IPL, and DO, while project administration was managed by AMA and DO. Funding acquisition was undertaken by DO.

## Acknowledgments

All circular detection analyses were conducted on the Hyperion cluster at the Donostia International Physics Center (DIPC). The author acknowledges the technical and human support provided by the DIPC Supercomputing Center. The authors have used generative AI technology (ChatGPT) to improve the readability and overall quality of the manuscript. The use of AI has not altered the content or messages conveyed in the manuscript, focusing solely on refining the writing for clarity and enhanced readability.

## Supplementary Information

**Figure S1.**
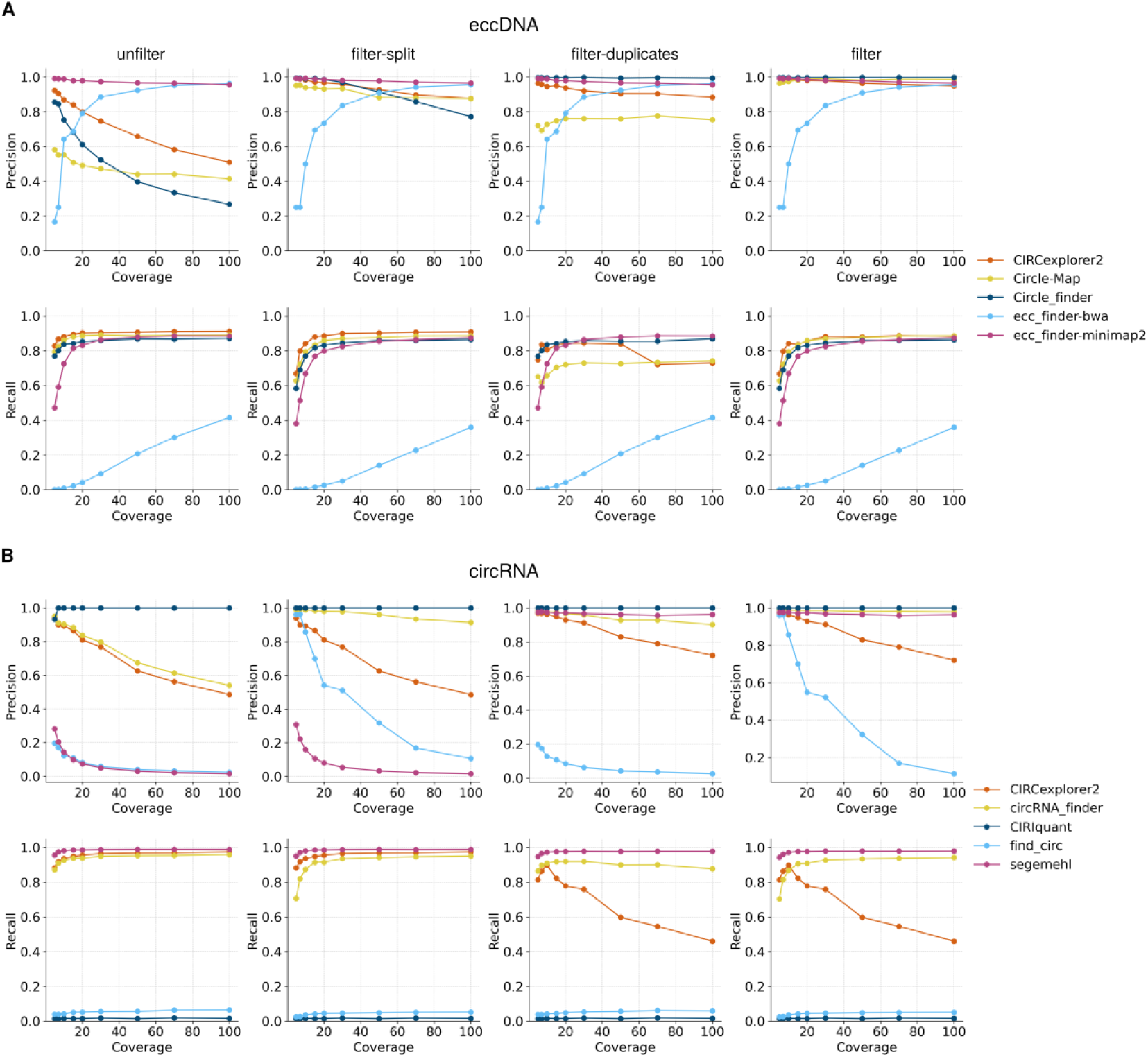
Performance analysis of detection software for eccDNA and circRNA identification in *in silico* datasets. Precision (**above**) and recall (**below**) values for eccDNA (**A**) and circRNA (**B**) detection across four filtering conditions: unfilter, filter-split, filter-duplicates, and filter. Color intensity indicates coverage level, with higher intensity corresponding to greater coverage.

**Figure S2.**
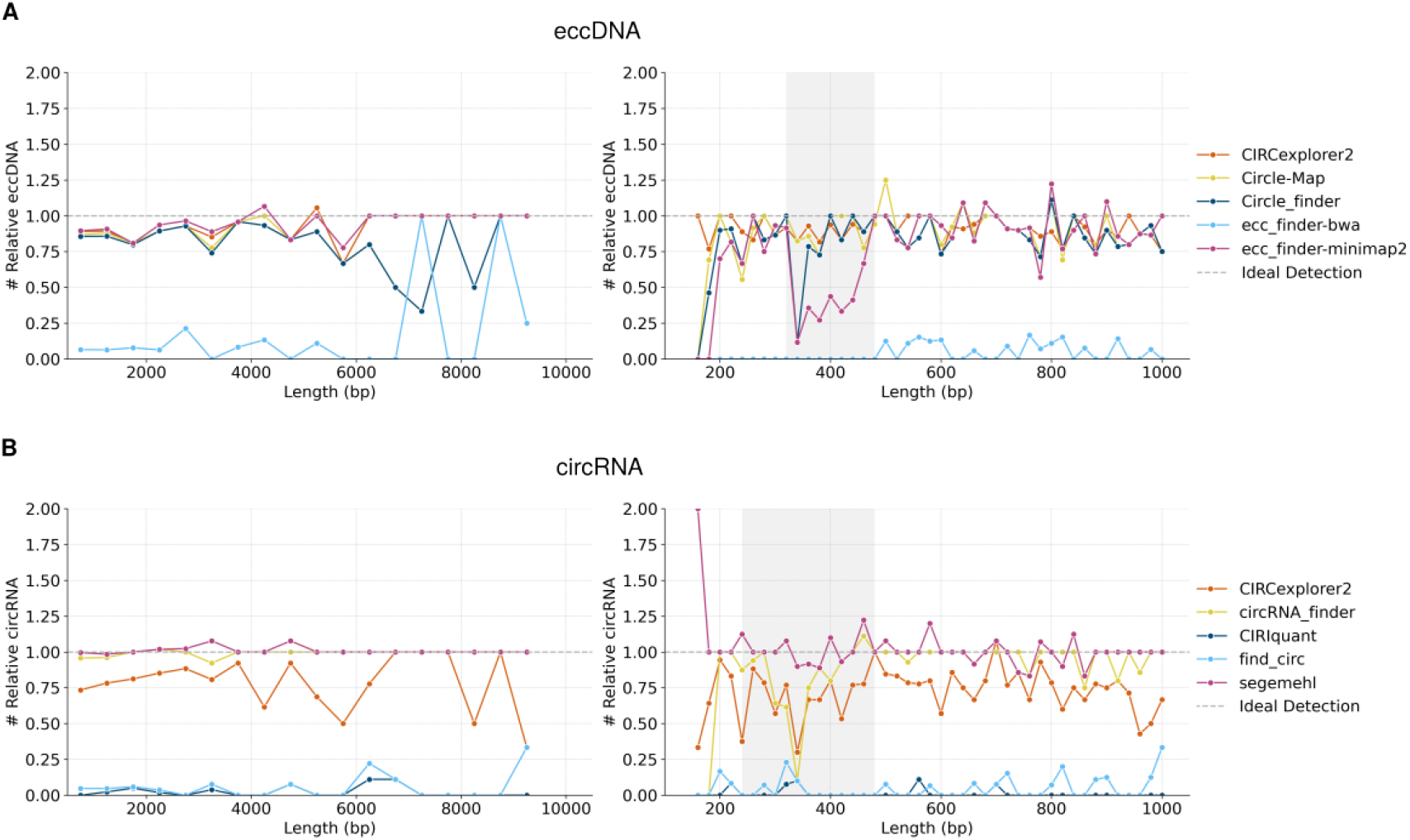
Circular length distribution analysis in *in silico* datasets. Relative circular length distribution of detected eccDNA (**A**) and circRNA (**B**) across all size ranges (**left**) and within the short-length range (**right**) in in silico datasets. A grey-shaded area highlights the length interval where detection performance was lowest (eccDNA: 320-480 bp; circRNA: 240-480 bp). To enhance the clarity of the distribution plots, a sliding window of size 5 was applied to smooth the distribution curves.

**Figure S3.**
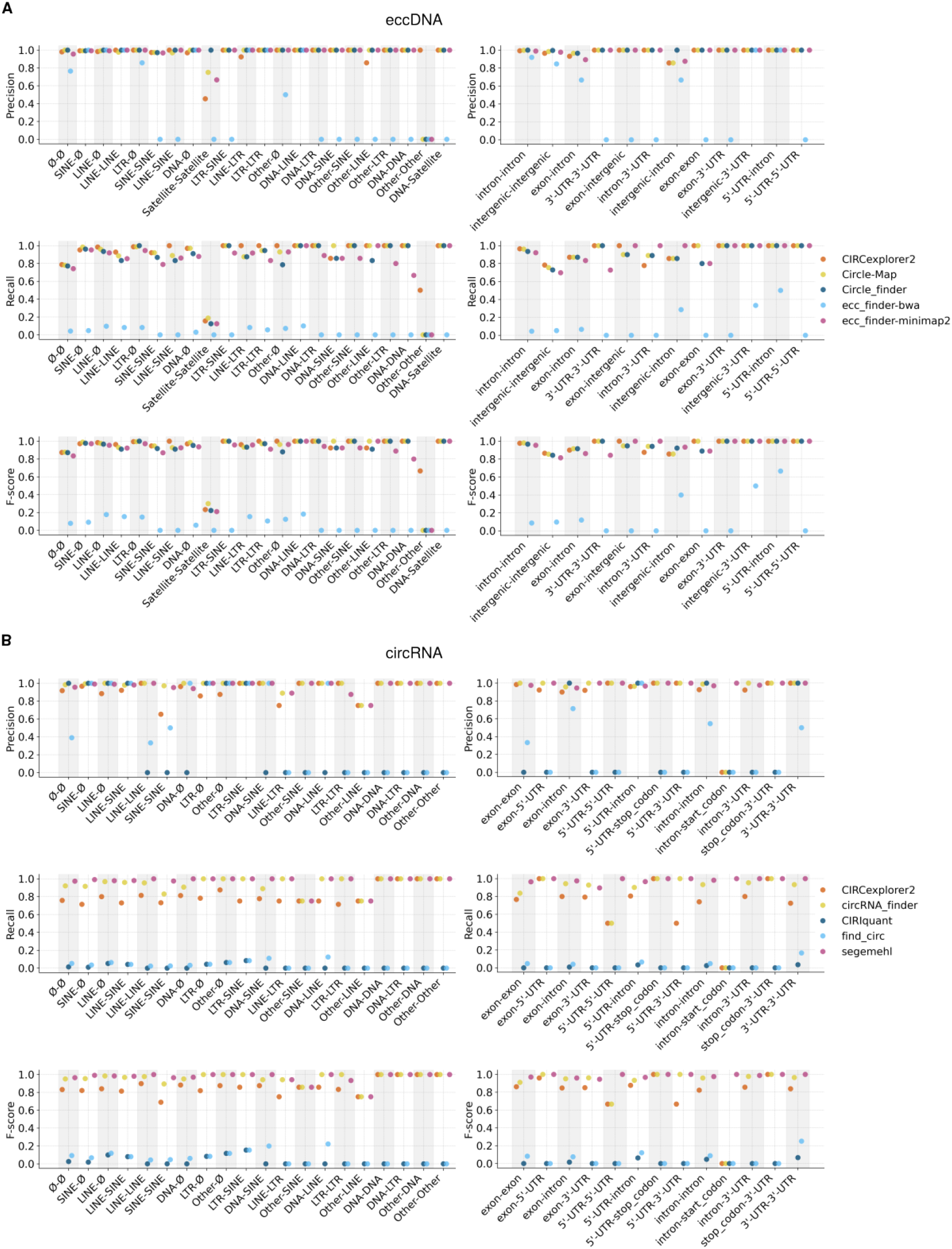
Repeat element analysis and genomic element analysis in *in silico* datasets. Precision (**above**), recall (**middle**) and F-score (**below**) values of all repeat elements (**left**) and genomic features (**right**) associated with detected eccDNA (**A**) and circRNA (**B**) in *in silico* datasets. In circRNAs no intergenic circles were generated and thus the region is not included in this plot.

**Figure S4.**
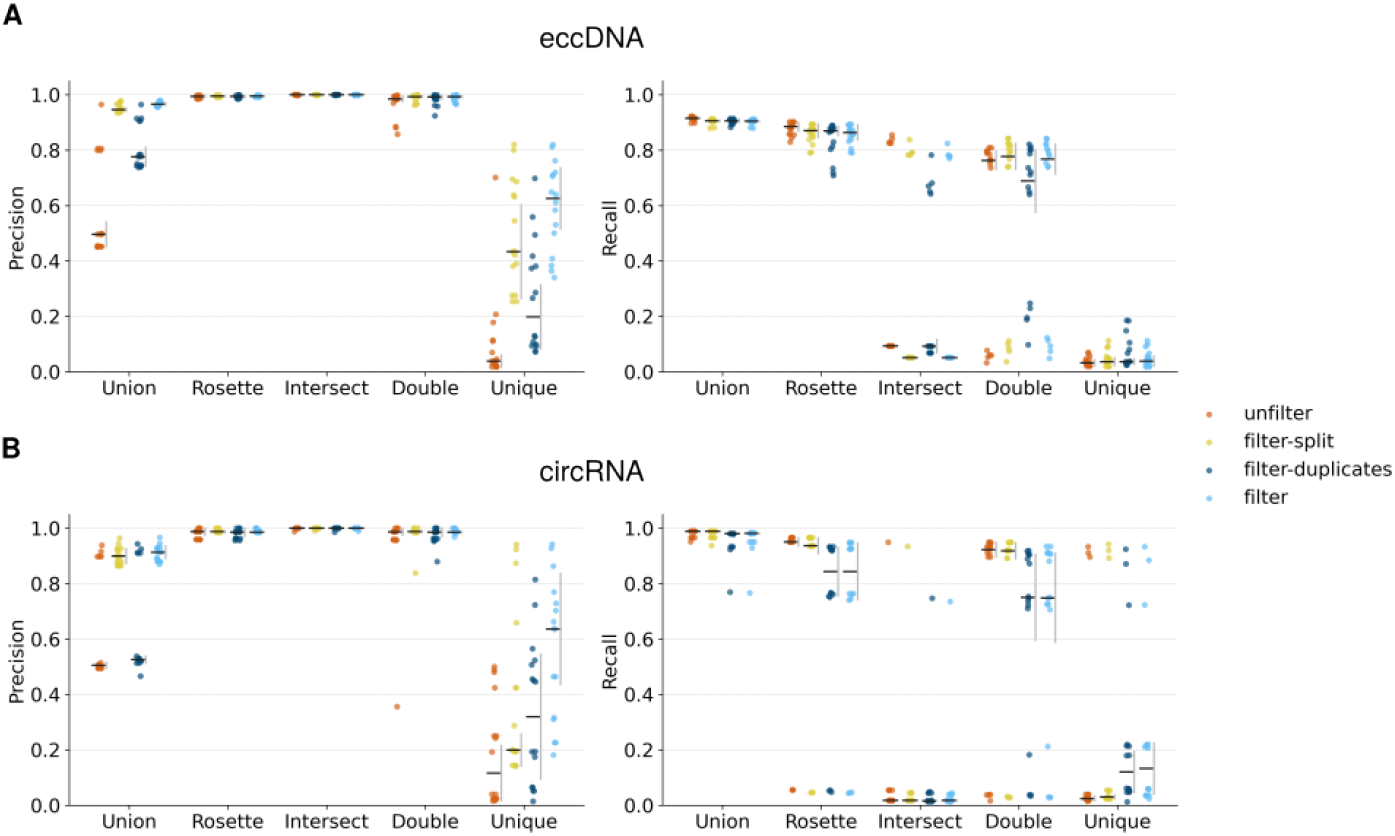
Performance analysis of software combinations for eccDNA and circRNA identification in *in silico* datasets. Strip plot of the F-score of software combination strategies—*Union, Rosette, Intersect, Double*, and *Unique*—were evaluated under four filtering conditions: *unfilter, filter-split, filter-duplicates*, and *filter*. Precision (**left**) and recall (**right**) values are shown for eccDNA (**A**) and circRNA (**B**). For each combination-filter pair, the horizontal bar represents the mean value and the vertical gray bar represents the standard deviation.

**Figure S5.**
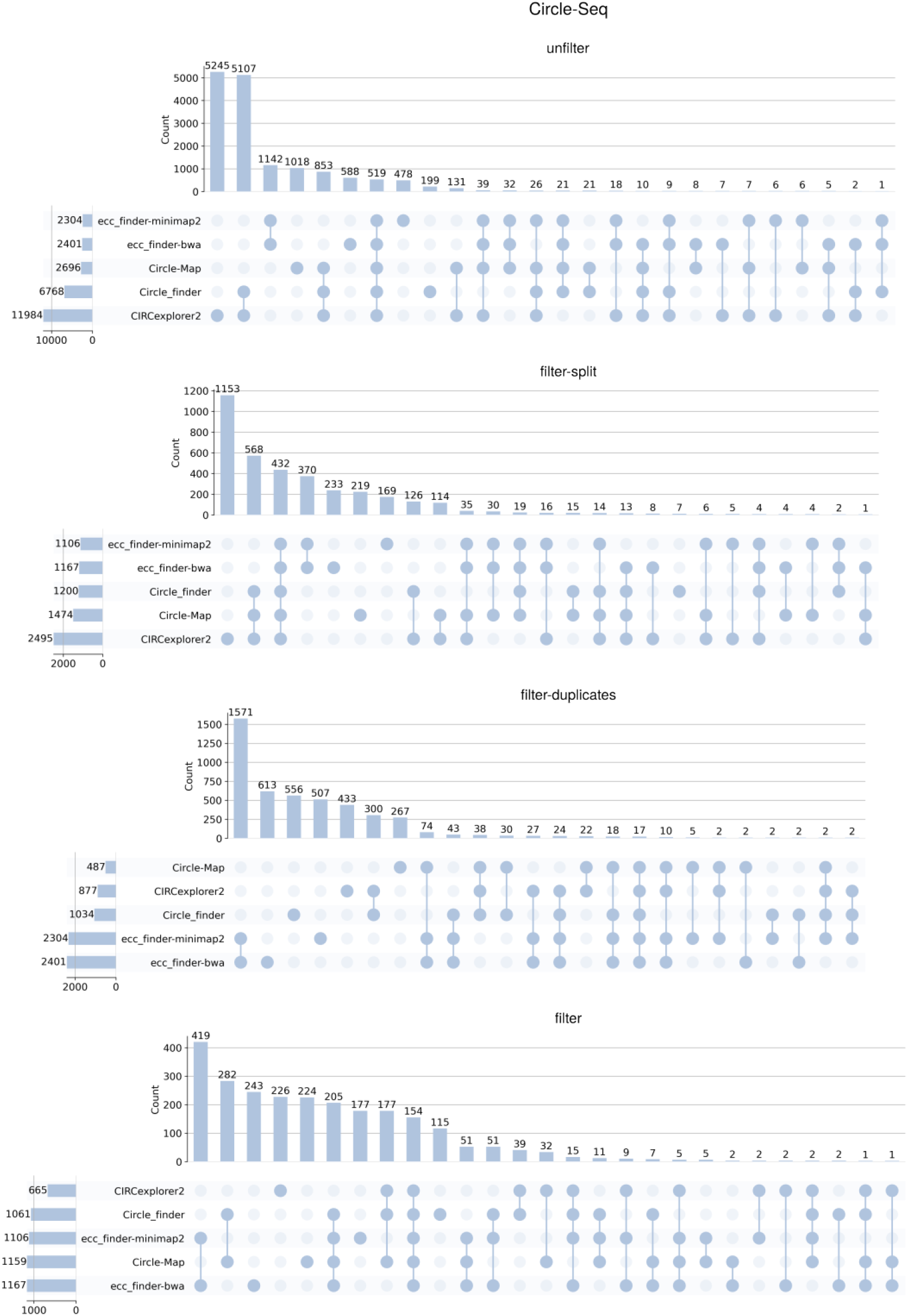
eccDNA detection in Circle-Seq data. UpSet plot showing the intersection of detected eccDNA among different detection tools in four filtering conditions: *unfilter, filter-split, filter-duplicates*, and *filter*. Each vertical bar represents the size of a specific intersection between tools, while the connected dots below indicate which tools are involved in each intersection. Horizontal bars on the left represent the total number of detections per tool.

**Figure S6.**
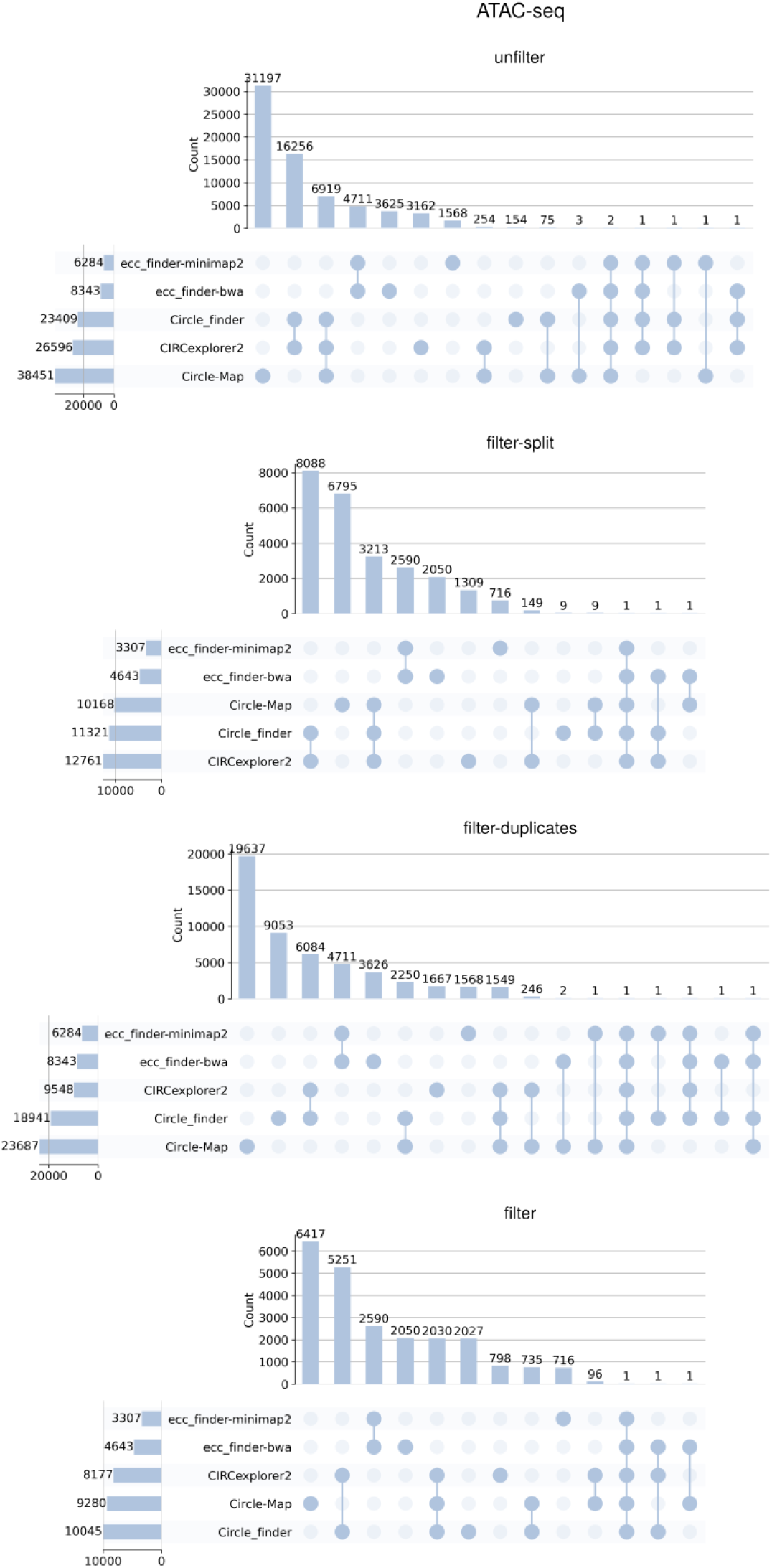
eccDNA detection in ATAC-seq data. UpSet plot showing the intersection of detected eccDNA among different detection tools in four filtering conditions: *unfilter*, filter-split, *filter-duplicates*, and *filter*. Each vertical bar represents the size of a specific intersection between tools, while the connected dots below indicate which tools are involved in each intersection. Horizontal bars on the left represent the total number of detections per tool.

**Figure S7.**
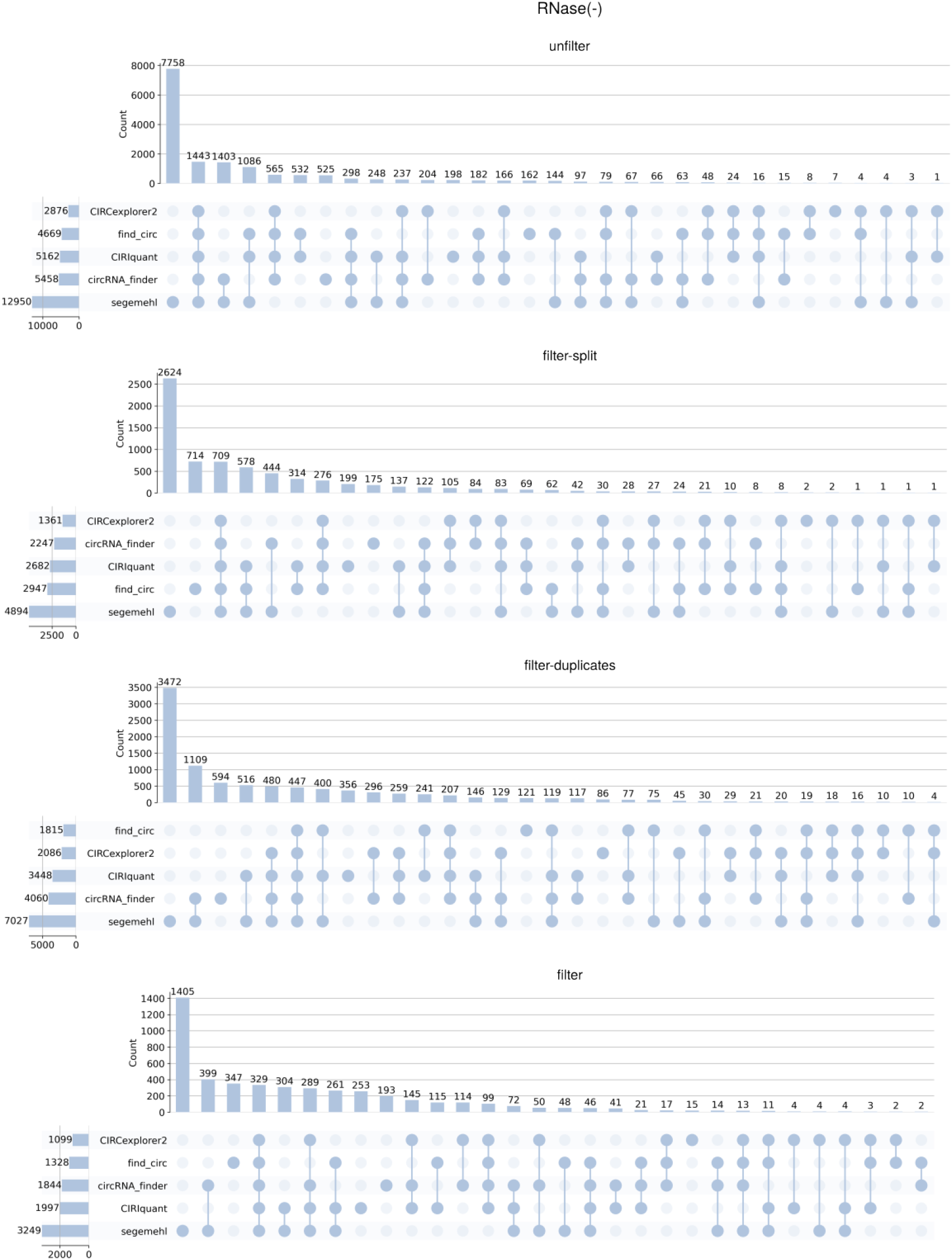
eccDNA detection in RNase(−) data. UpSet plot showing the intersection of detected circRNA among different detection tools in four filtering conditions: *unfilter, filter-split, filter-duplicates*, and *filter*. Each vertical bar represents the size of a specific intersection between tools, while the connected dots below indicate which tools are involved in each intersection. Horizontal bars on the left represent the total number of detections per tool.

**Figure S8.**
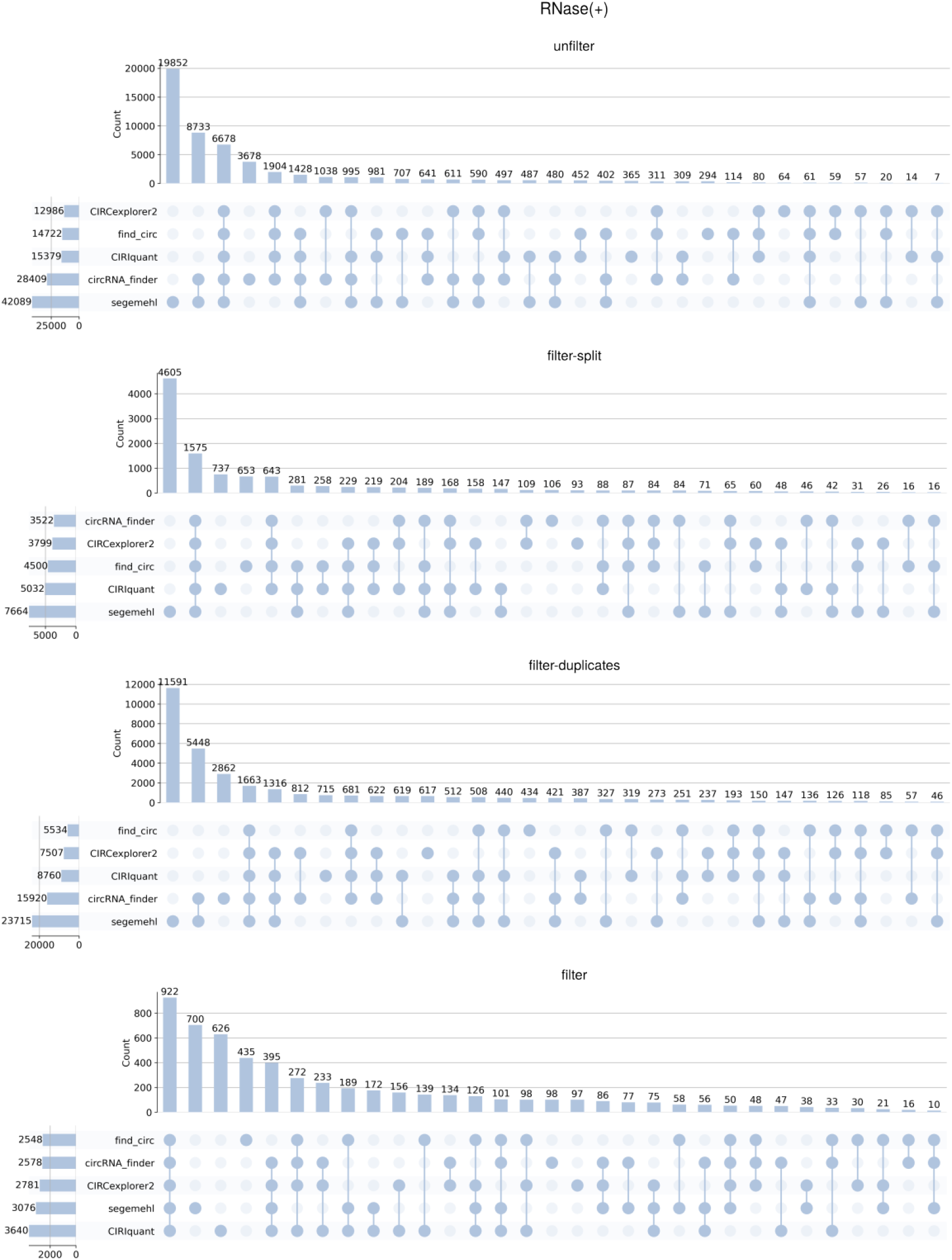
circRNA detection in RNase(+) data. UpSet plot showing the intersection of detected circRNA among different detection tools in four filtering conditions: *unfilter, filter-split, filter-duplicates*, and *filter*. Each vertical bar represents the size of a specific intersection between tools, while the connected dots below indicate which tools are involved in each intersection. Horizontal bars on the left represent the total number of detections per tool.

**Figure S9.**
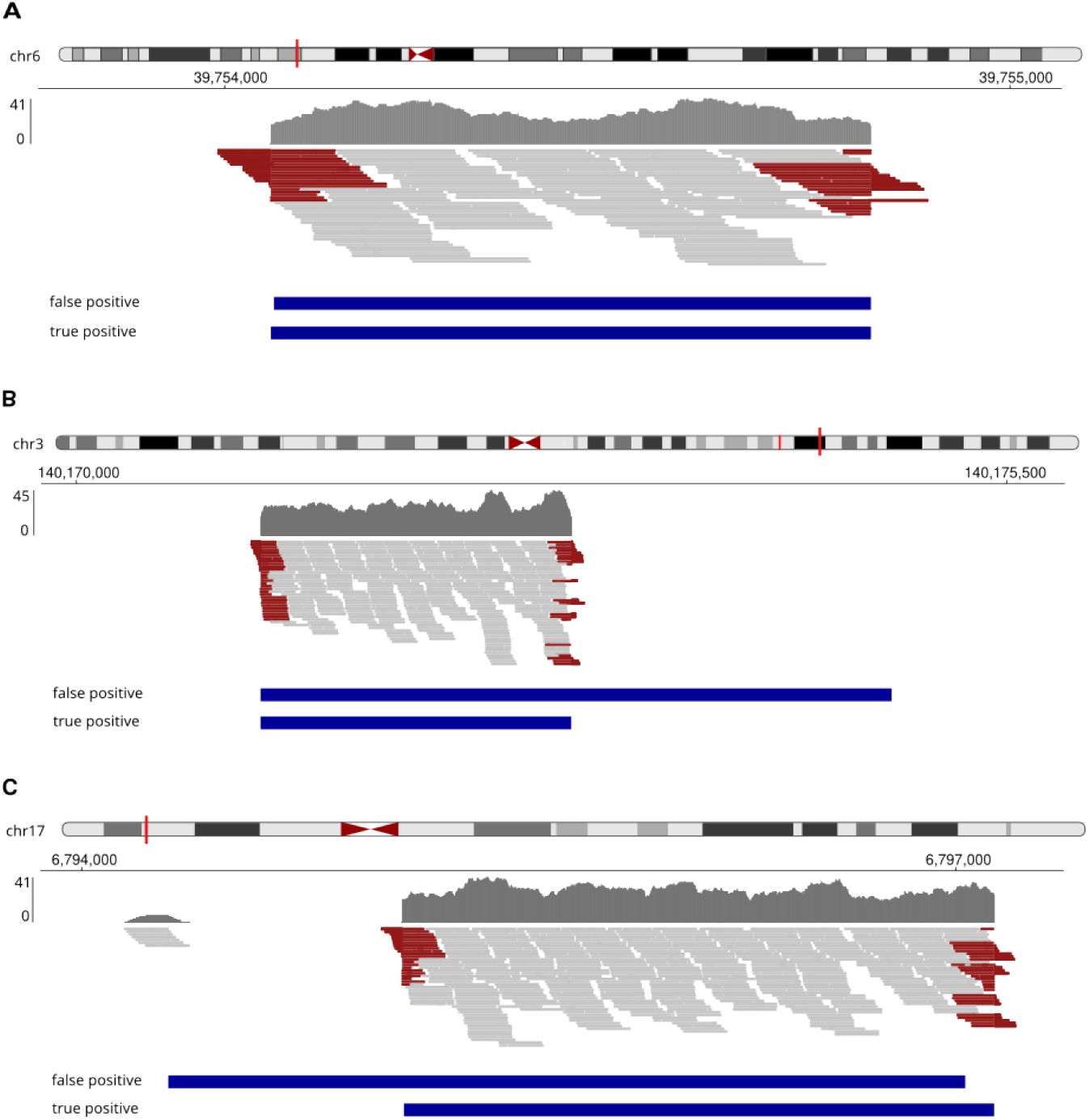
IGV visualization of three eccDNA false positives in *in silico* datasets. Genome browser views (IGV) illustrating three examples of false positive (FP) eccDNAs, each shown alongside their corresponding true positive (TP) circle. (**A**) TP: chr6:39,754,059–39,754,823; FP: chr6:39,754,063–39,754,823. (**B**) TP: chr3:140,171,091–140,172,928; FP: chr3:140,171,091–140,174,822. (**C**) TP: chr3:140,171,091–140,172,928; FP: chr17:6,794,297–6,797,033. Reads marked in red represent the split reads associated with the CJ.

**Figure S10.**
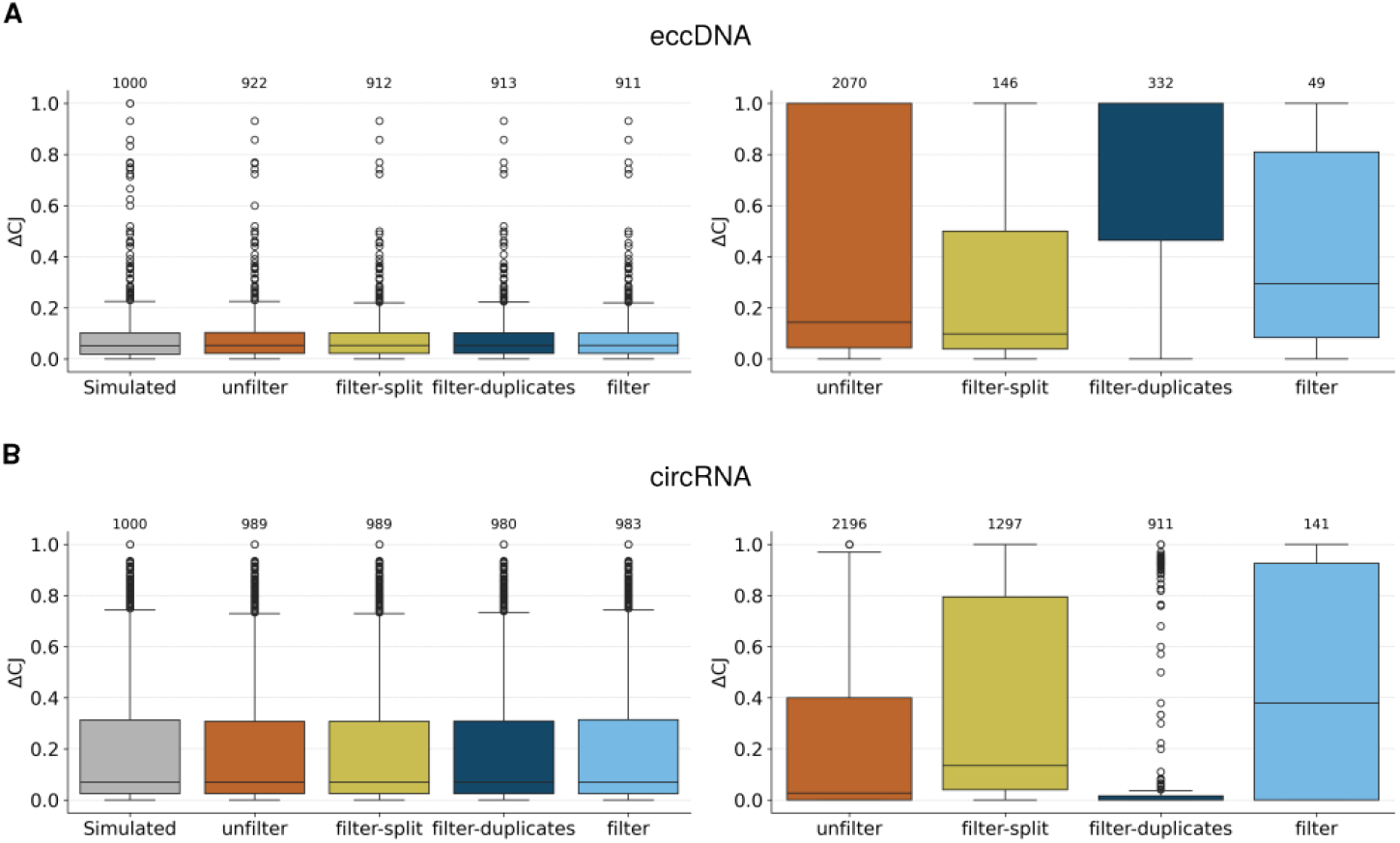
Circular junction nucleotide difference (ΔCJ) in *in silico* datasets. Boxplots of ΔCJ values of TP (**left**) and FP (**right**) circles for (**A**) eccDNA and (**B**) circRNA under *filter* filtering.

**Figure S11.**
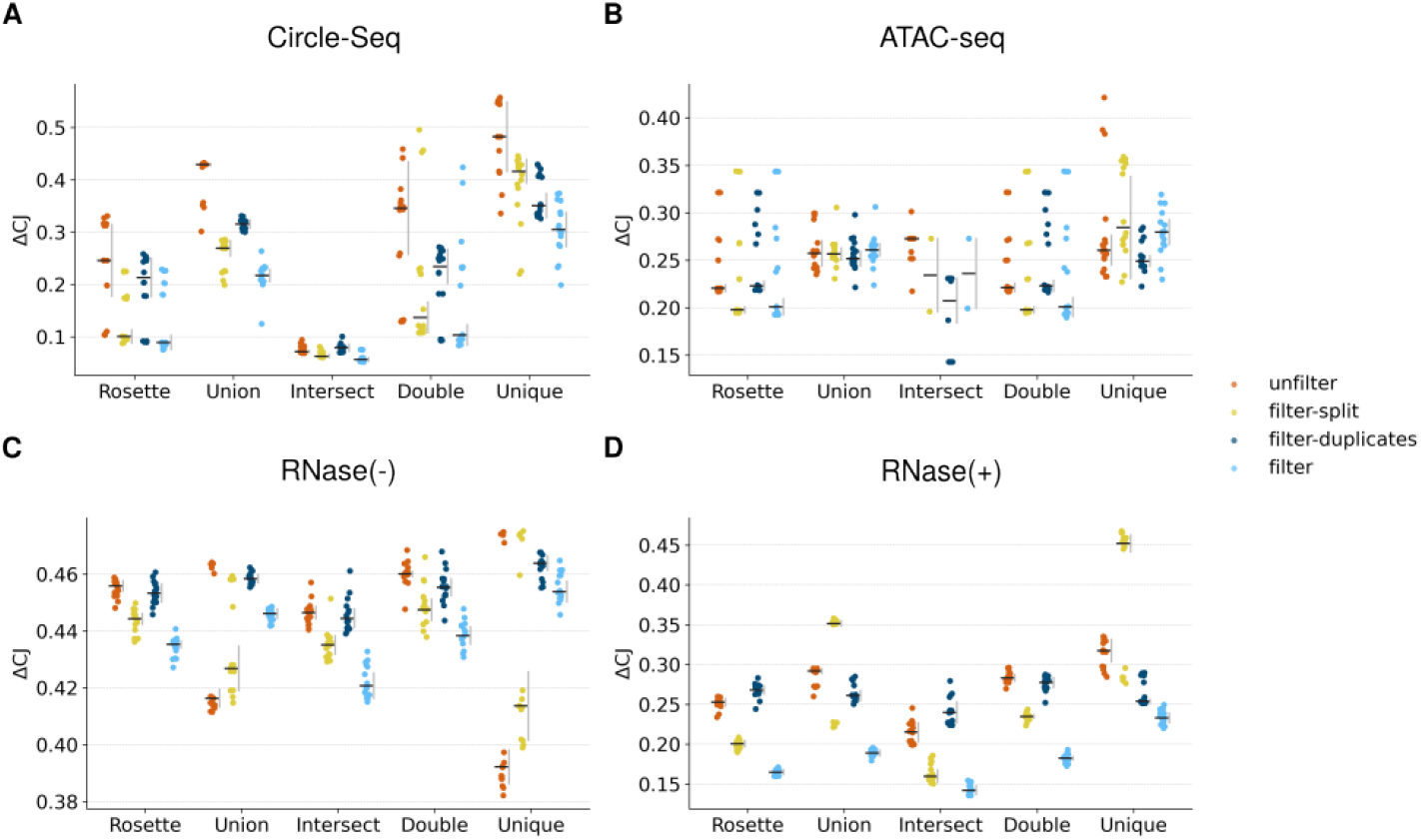
Circular junction nucleotide difference (ΔCJ) of software combinations for eccDNA and circRNA identification in biological datasets. Strip plot of ΔCJ values for different tool combinations—Union, Rosette, Intersect, Double, and Unique—evaluated on biological data. Results are shown for Circle-Seq (**A**) and ATAC-seq (**B**) data for eccDNA, and RNase(−) (**C**) and RNase(+) (**D**) data for circRNA, under four filtering conditions: *unfilter, filter-split, filter-duplicates*, and *filter*.

**Figure S12.**
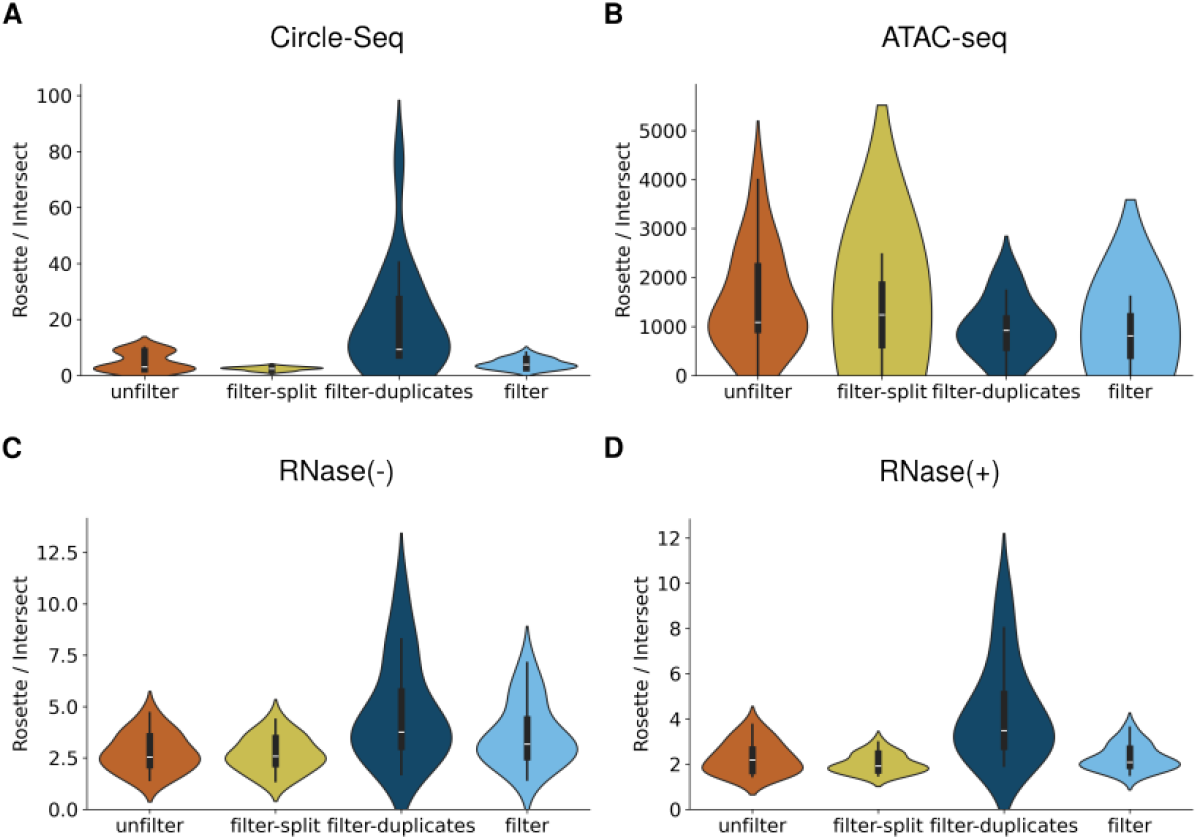
Comparison of detected circles between Rosette and Intersect. Violin plots of the ratio of detected circles in different tool combinations between *Rosette* and *Intersect*in biological data. Results are shown for Circle-Seq (**A**) and ATAC-seq data (**B**) for eccDNA, and RNase(−) (**C**) and RNase(+) (**D**) data for circRNA, under four filtering conditions: *unfilter, filter-split, filter-duplicates*, and *filter*.

**Figure S13.**
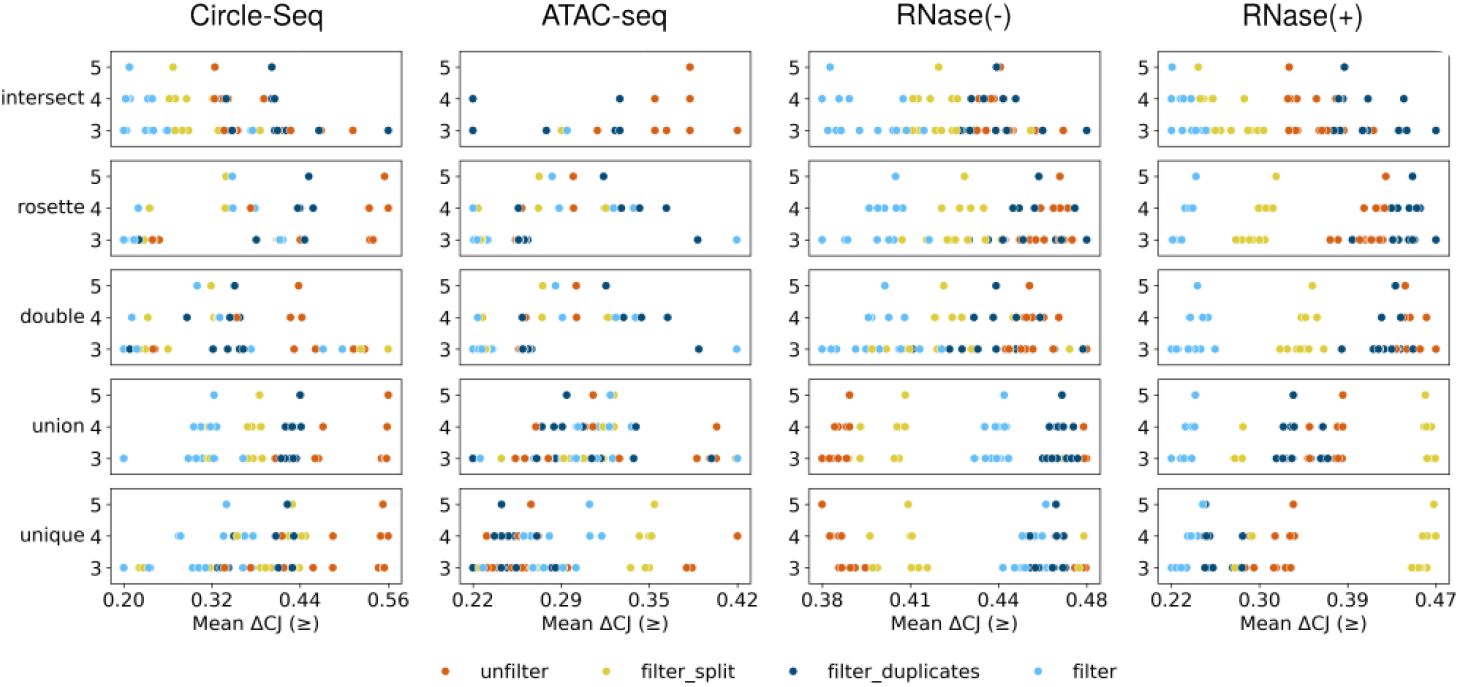
Performance of tool combinations for eccDNA and circRNA in biological datasets. Circular junction nucleotide difference (ΔCJ) mean values are reported, indicating the number of tools combined, across different combinations, filtering conditions and circle enrichment strategies.

### Additional files

**Supplementary Material 1**. Performance analysis of detection software for eccDNA identification in *in silico* datasets.

**Supplementary Material 2**. Performance analysis of detection software for circRNA identification in *in silico* datasets.

**Supplementary Material 3**. Kolmogorov-Smirnov test for circular length distribution in *in silico* datasets.

**Supplementary Material 4**. Repeat element annotation for eccDNA in *in silico* datasets.

**Supplementary Material 5**. Repeat element annotation for circRNA in *in silico* datasets.

**Supplementary Material 6**. Genomic element annotation for eccDNA in *in silico* datasets.

**Supplementary Material 7**. Genomic element annotation for circRNA in *in silico* datasets.

**Supplementary Material 8**. Performance analysis of software combinations detection for eccDNA in *in silico unfilter* condition datasets.

**Supplementary Material 9**. Performance analysis of software combinations detection for eccDNA in *in silico filter-split* condition datasets.

**Supplementary Material 10**. Performance analysis of software combinations detection for eccDNA in *in silico filter-duplicates* condition datasets.

**Supplementary Material 11**. Performance analysis of software combinations detection for eccDNA in *in silico filter* condition datasets.

**Supplementary Material 12**. Dunn’s test results for software combination detection performance metrics for eccDNA in *in silico* datasets.

**Supplementary Material 13**. Performance analysis of software combinations detection for circRNA in *in silico unfilter* condition datasets.

**Supplementary Material 14**. Performance analysis of software combinations detection for circRNA in *in silico filter-split* condition datasets.

**Supplementary Material 15**. Performance analysis of software combinations detection for circRNA in *in silico filter-duplicates* condition datasets.

**Supplementary Material 16**. Performance analysis of software combinations detection for circRNA in *in silico filter* condition datasets.

**Supplementary Material 17**. Dunn’s test results for software combination detection performance metrics for circRNA in *in silico* datasets.

**Supplementary Material 18**. Circular detection in biological datasets.

**Supplementary Material 19**. ΔCJ evaluation as a proxy measure of circle detection quality for eccDNA identification in *in silico* datasets.

**Supplementary Material 20**. ΔCJ evaluation as a proxy measure of circle detection quality for circRNA identification in *in silico* datasets.

**Supplementary Material 21**. Performance analysis of detection software in biological datasets.

**Supplementary Material 22**. Performance analysis of detection software combinations in biological datasets.

**Supplementary Material 23**. Dunn’s test results for software combination detection performance metrics in biological datasets.

## Bibliography

Chen, L.-L. and Yang, L. (2015). Regulation of circrna biogenesis. RNA biology, 12(4):381–388.

Cohen, S. and Mechali, M. (2001). A novel cell-free system reveals a mechanism of circular dna formation from tandem repeats. Nucleic acids research, 29(12):2542–2548.

Di Tommaso, P., Chatzou, M., Floden, E. W., Barja, P. P., Palumbo, E., and Notredame, C. (2017). Nextflow enables reproducible computational workflows. Nature biotechnology, 35(4):316–319.

Digby, B., Finn, S. P., and Ó Broin, P. (2023). nf-core/circrna: a portable workflow for the quantification, mirna target prediction and differential expression analysis of circular rnas. BMC bioinformatics, 24(1):27.

Digby, B., Finn, S., and Ó Broin, P. (2024). Computational approaches and challenges in the analysis of circrna data. BMC genomics, 25(1):527.

Dobin, A., Davis, C. A., Schlesinger, F., Drenkow, J., Zaleski, C., Jha, S., Batut, P., Chaisson, M., and Gingeras, T. R. (2013). Star: ultrafast universal rna-seq aligner. Bioinformatics, 29(1):15–21.

Ewels, P. A., Peltzer, A., Fillinger, S., Patel, H., Alneberg, J., Wilm, A., Garcia, M. U., Di Tommaso, P., and Nahnsen, S. (2020). The nf-core framework for community-curated bioinformatics pipelines. Nature biotechnology, 38(3):276–278.

Gaffo, E., Buratin, A., Dal Molin, A., and Bortoluzzi, S. (2021). Sensitive, reliable and robust circrna detection from rna-seq with circompara2. Briefings in Bioinformatics, 23(1). doi: 10.1093/bib/bbab418.

Gao, X., Liu, K., Luo, S., Tang, M., Liu, N., Jiang, C., Fang, J., Li, S., Hou, Y., Guo, C., et al. (2024). Comparative analysis of methodologies for detecting extrachromosomal circular dna. Nature Communications, 15(1):9208.

Gao, Y., Wang, J., and Zhao, F. (2015). Ciri: an efficient and unbiased algorithm for de novo circular rna identification. Genome biology, 16:1–16.

Hansen, T. B. (2018). Improved circrna identification by combining prediction algorithms. Frontiers in cell and developmental biology, 6:330528.

Hoffmann, S., Otto, C., Kurtz, S., Sharma, C. M., Khaitovich, P., Vogel, J., Stadler, P. F., and Hackermüller, J. (2009). Fast mapping of short sequences with mismatches, insertions and deletions using index structures. PLoS computational biology, 5 (9):e1000502.

Iparraguirre, L., Muñoz-Culla, M., Prada-Luengo, I., Castillo-Triviño, T., Olascoaga, J., and Otaegui, D. (2017). Circular rna profiling reveals that circular rnas from anxa2 can be used as new biomarkers for multiple sclerosis. Human molecular genetics, 26(18):3564–3572.

Iparraguirre, L., Prada-Luengo, I., Regenberg, B., and Otaegui, D. (2019). To be or not to be: circular rnas or mrnas from circular dnas? Frontiers in Genetics, 10:940.

Jeck, W. R., Sorrentino, J. A., Wang, K., Slevin, M. K., Burd, C. E., Liu, J., Marzluff, W. F., and Sharpless, N. E. (2013). Circular rnas are abundant, conserved, and associated with alu repeats. Rna, 19(2):141–157.

Kim, D., Pertea, G., Trapnell, C., Pimentel, H., Kelley, R., and Salzberg, S. L. (2013). Tophat2: accurate alignment of transcriptomes in the presence of insertions, deletions and gene fusions. Genome biology, 14:1–13.

Kumar, P., Dillon, L. W., Shibata, Y., Jazaeri, A. A., Jones, D. R., and Dutta, A. (2017). Normal and cancerous tissues release extrachromosomal circular dna (eccdna) into the circulation. Molecular Cancer Research, 15(9):1197–1205.

Kumar, P., Kiran, S., Saha, S., Su, Z., Paulsen, T., Chatrath, A., Shibata, Y., Shibata, E., and Dutta, A. (2020). Atac-seq identifies thousands of extrachromosomal circular dna in cancer and cell lines. Science advances, 6(20):eaba2489.

Lex, A., Gehlenborg, N., Strobelt, H., Vuillemot, R., and Pfister, H. (2014). Upset: Visualization of intersecting sets. IEEE Transactions on Visualization and Computer Graphics, 20(12):1983–1992. doi: 10.1109/TVCG.2014.2346248.

Li, F., Ming, W., Lu, W., Wang, Y., Dong, X., and Bai, Y. (2024). Bioinformatics advances in eccdna identification and analysis. Oncogene, 43(41):3021–3036.

Li, H. (2018). Minimap2: pairwise alignment for nucleotide sequences. Bioinformatics, 34(18):3094–3100.

Li, H. and Durbin, R. (2009). Fast and accurate short read alignment with burrows–wheeler transform. bioinformatics, 25(14): 1754–1760.

Liu, H., Akhatayeva, Z., Pan, C., Liao, M., and Lan, X. (2022). Comprehensive comparison of two types of algorithm for circrna detection from short-read rna-seq. Bioinformatics, 38(11):3037–3043. doi: 10.1093/bioinformatics/btac302.

Memczak, S., Jens, M., Elefsinioti, A., Torti, F., Krueger, J., Rybak, A., Maier, L., Mackowiak, S. D., Gregersen, L. H., Munschauer, M., et al. (2013). Circular rnas are a large class of animal rnas with regulatory potency. Nature, 495(7441):333–338.

Møller, H. D., Parsons, L., Jørgensen, T. S., Botstein, D., and Regenberg, B. (2015). Extrachromosomal circular dna is common in yeast. Proceedings of the National Academy of Sciences, 112(24):E3114–E3122.

Møller, H. D., Bojsen, R. K., Tachibana, C., Parsons, L., Botstein, D., and Regenberg, B. (2016). Genome-wide purification of extrachromosomal circular dna from eukaryotic cells. JoVE (Journal of Visualized Experiments), (110):e54239.

Møller, H. D., Mohiyuddin, M., Prada-Luengo, I., Sailani, M. R., Halling, J. F., Plomgaard, P., Maretty, L., Hansen, A. J., Snyder, M. P., Pilegaard, H., et al. (2018). Circular dna elements of chromosomal origin are common in healthy human somatic tissue. Nature communications, 9(1):1069.

Møller, H. D., Ramos-Madrigal, J., Prada-Luengo, I., Gilbert, M. T. P., and Regenberg, B. (2020). Near-random distribution of chromosome-derived circular dna in the condensed genome of pigeons and the larger, more repeat-rich human genome. Genome biology and evolution, 12(2):3762–3777.

Noer, J. B., Hørsdal, O. K., Xiang, X., Luo, Y., and Regenberg, B. (2022). Extrachromosomal circular dna in cancer: history, current knowledge, and methods. Trends in Genetics, 38(7):766–781. doi: 10.1016/j.tig.2022.02.007.

Prada-Luengo, I., Krogh, A., Maretty, L., and Regenberg, B. (2019). Sensitive detection of circular dnas at single-nucleotide resolution using guided realignment of partially aligned reads. BMC bioinformatics, 20:1–9.

Salzman, J., Gawad, C., Wang, P. L., Lacayo, N., and Brown, P. O. (2012). Circular rnas are the predominant transcript isoform from hundreds of human genes in diverse cell types. PloS one, 7(2):e30733.

Sanger, H. L., Klotz, G., Riesner, D., Gross, H. J., and Kleinschmidt, A. K. (1976). Viroids are single-stranded covalently closed circular rna molecules existing as highly base-paired rod-like structures. Proceedings of the National Academy of Sciences, 73(11):3852–3856.

Schreyer, D., nf-core bot, Ewels, P., and Peltzer, A. nf-core/circdna: v1.1.0 – tremendous wombat, (2024). URL 10.5281/zenodo.10643212.

Sin, S. T., Deng, J., Ji, L., Yukawa, M., Chan, R. W., Volpi, S., Vaglio, A., Fenaroli, P., Bocca, P., Cheng, S. H., et al. (2022). Effects of nucleases on cell-free extrachromosomal circular dna. JCI insight, 7(8).

Szabo, L. and Salzman, J. (2016). Detecting circular rnas: bioinformatic and experimental challenges. Nature Reviews Genetics, 17(11):679–692.

Tsai, S. Q., Nguyen, N. T., Malagon-Lopez, J., Topkar, V. V., Aryee, M. J., and Joung, J. K. (2017). Circle-seq: a highly sensitive in vitro screen for genome-wide crispr–cas9 nuclease off-targets. Nature Methods, 14(6):607–614. doi: 10.1038/nmeth.4278.

Turner, K. M., Deshpande, V., Beyter, D., Koga, T., Rusert, J., Lee, C., Li, B., Arden, K., Ren, B., Nathanson, D. A., et al. (2017). Extrachromosomal oncogene amplification drives tumour evolution and genetic heterogeneity. Nature, 543(7643):122–125.

Vromman, M., Anckaert, J., Bortoluzzi, S., Buratin, A., Chen, C.-Y., Chu, Q., Chuang, T.-J., Dehghannasiri, R., Dieterich, C., Dong, X., et al. (2023). Large-scale benchmarking of circrna detection tools reveals large differences in sensitivity but not in precision. Nature methods, 20(8):1159–1169.

Westholm, J. O., Miura, P., Olson, S., Shenker, S., Joseph, B., Sanfilippo, P., Celniker, S. E., Graveley, B. R., and Lai, E. C. (2014). Genome-wide analysis of drosophila circular rnas reveals their structural and sequence properties and age-dependent neural accumulation. Cell reports, 9(5):1966–1980.

Yu, J., Zhang, H., Han, P., Jiang, X., Li, J., Li, B., Yang, S., He, C., Mao, S., Dang, Y., and Xiang, X. (2024). Circle-seq based method for eccdna synthesis and its application as a canonical promoter independent vector for robust microrna overexpression. Computational and Structural Biotechnology Journal, 23:358–368. doi: 10.1016/j.csbj.2023.12.019.

Zeng, X., Lin, W., Guo, M., and Zou, Q. (2017). A comprehensive overview and evaluation of circular rna detection tools. PLoS computational biology, 13(6):e1005420.

Zhang, H.-d., Jiang, L.-h., Sun, D.-w., Hou, J.-c., and Ji, Z.-l. (2018). Circrna: a novel type of biomarker for cancer. Breast cancer, 25:1–7.

Zhang, J., Chen, S., Yang, J., and Zhao, F. (2020). Accurate quantification of circular rnas identifies extensive circular isoform switching events. Nature communications, 11(1):90.

Zhang, P., Peng, H., Llauro, C., Bucher, E., and Mirouze, M. (2021). ecc_finder: a robust and accurate tool for detecting extrachromosomal circular dna from sequencing data. Frontiers in plant science, 12:743742.

Zhang, X.-O., Wang, H.-B., Zhang, Y., Lu, X., Chen, L.-L., and Yang, L. (2014). Complementary sequence-mediated exon circularization. Cell, 159(1):134–147.

Zhang, X.-O., Dong, R., Zhang, Y., Zhang, J.-L., Luo, Z., Zhang, J., Chen, L.-L., and Yang, L. (2016). Diverse alternative back-splicing and alternative splicing landscape of circular rnas. Genome research, 26(9):1277–1287.

Zhao, Y., Yu, L., Zhang, S., Su, X., and Zhou, X. (2022). Extrachromosomal circular dna: Current status and future prospects. eLife, 11. doi: 10.7554/elife.81412.

